# Wheat diversity reveals new genomic loci and candidate genes for vegetation indices using genome-wide association analysis

**DOI:** 10.64898/2026.01.14.699455

**Authors:** Samira Rustamova, Atabay Jahangirov, Jens Léon, Ali Ahmad Naz, Irada Huseynova

**Affiliations:** Institute of Molecular Biology of the Ministry of Science and Education of the Republic of Azerbaijan, Baku, Azerbaijan; Research Institute of Crop Husbandry of the Ministry of Agriculture of the Republic of Azerbaijan, Gobustan, Azerbaijan; Field Lab Campus Klein-Altendorf, University of Bonn, Rheinbach, Germany; Department of Plant Breeding, University of Applied Sciences, Osnabrueck, Germany; Department of Molecular Biology and Biotechnologies, Baku State University, Baku, Azerbaijan

**Keywords:** Genetic diversity, Digital phenotyping, Heritability, GWAS, Quantitative trait loci, Marker-assisted selection

## Abstract

Wheat (*Triticum aestivum* L.), a globally essential crop, exhibits high vulnerability to drought stress, particularly within rainfed agricultural systems. Enhancing resilience requires a deeper understanding of the genetic architecture underlying key physiological traits. Spectral vegetation indices provide a high-throughput, non-invasive approach to quantify these traits, but their genetic basis in wheat under drought remains largely unexplored. We conducted a genome-wide association study (GWAS) using 187 bread wheat genotypes evolved and selected across rainfed conditions. This population was phenotyped for 25 vegetation indices and genotyped using a 25K SNP array. Phenotypic data showed significant genetic variation with broad-sense heritability (*H*²) ranging from 0.19 to 0.95. Comparing phenotype and genotype data identified 812 Bonferroni-significant associations distributed across the A, B and D genomes. A prominent major QTL effect was identified as a hotspot on chromosome 2A, tagged by SNP marker wsnp_Ex_c36049_44083089, was among the strongest associations for 17 vegetation indices, explaining up to 20% of the genotypic variance for key traits like greenness and pigment indices. Candidate gene analysis at this locus identified the co-localization of a *LEA_2/NDR1-like* gene and a lectin receptor-like kinase with multiple genes involved in terpenoid, phenylpropanoid and primary metabolism, consistent with integrated roles in stress signaling and metabolic acclimation. These data show that vegetation indices are heritable digital phenotypes, which can be employed for the selection and genetic analysis of essential physiological growth parameters under varying and adverse climatic conditions.

## Introduction

Global food security is intrinsically linked to the productivity of wheat, a crop that supplies approximately 18% of dietary energy and 19% of protein worldwide (FAO et al. 2025). According to recent projections, global agricultural markets are facing a precarious supply-demand balance, where decelerating yield growth rates are increasingly struggling to keep pace with the sustained expansion of consumption (OECD/FAO 2023). The stability of wheat production is increasingly threatened by climate change, which intensifies abiotic stresses, particularly drought and extreme heat, and thereby threatens the sustainability of food systems (IPCC 2022). As a result, rainfed production systems, which account for a significant share of global wheat supply, face disproportionately high and rising risks, particularly in semi-arid and arid regions (FAO 2020). Empirical studies show that grain yield losses can exceed 40% under terminal drought and can be even greater if water stress coincides with heat stress during the reproductive phases (Qaseem et al. 2019a, b).

For the South Caucasus, and Azerbaijan in particular, drought poses a strategic challenge. Current climate assessments project a sustained temperature increase of 1–3 °C by approximately 2050, coupled with a reduction in annual precipitation, especially during the summer months, which will directly affect the productivity of rainfed crops (Red Cross fact sheet 2024). More than 40% of the national wheat area relies solely on natural precipitation, making production highly vulnerable to climatic variability (Talai et al. 2023). Such trends represent a serious challenge for national food security, as wheat constitutes the major staple in the local diet and is a strategic crop for ensuring self-sufficiency. Under these conditions, enhancing the resilience of wheat germplasm to water-limited environments is of utmost importance for stabilizing yields and supporting sustainable agriculture.

Drought tolerance is one of the most complex agronomic traits, being polygenic, characterized by low heritability, and strongly influenced by genotype × environment interactions (Bapela et al. 2022). This complexity is further amplified in bread wheat (*Triticum aestivum* L.) due to the challenges of trait dissection in its large, highly redundant hexaploid genome (Borrill et al. 2019). Advances in genomics, such as high-density single-nucleotide polymorphism (SNP) arrays and genotyping-by-sequencing (GBS), have enabled high-resolution mapping of quantitative trait loci (QTL) for complex traits, including stress resilience, yield, and grain quality, via genome-wide association studies (GWAS) (Sharma et al. 2025).Unlike traditional biparental linkage mapping, GWAS leverages historical recombination events within diverse germplasm panels to identify marker-trait associations (MTAs) with higher resolution, offering greater relevance to breeding. The release of the IWGSC reference genome (IWGSC 2018) has been pivotal, providing a robust framework for anchoring SNPs to physical positions and pinpointing candidate genes.

In wheat, GWAS has been successfully applied to dissect the genetic architecture of key agronomic traits. For instance, this approach has identified stable QTLs for grain yield and its components under drought and heat stress in the CIMMYT durum wheat panel (Sukumaran et al. 2018), as well as for radiation use efficiency and biomass accumulation in spring wheat (Molero et al. 2019). GWAS has also enabled the identification of loci associated with heading date variation under autumn-sowing conditions (Kim et al. 2025), loci controlling yield-related traits in bread wheat under combined pre-anthesis drought and heat stress (Qaseem et al. 2019b), and grain yield and quality under contrasting water regimes (Govta et al. 2022). This method has been extended to physiological and biochemical markers, such as grain peroxidase activity (Zhou et al. 2021), calcium accumulation in grains (Shi et al., 2022), drought-induced proline accumulation (Kamruzzaman et al. 2023), and hydrogen peroxide response in root and shoot tissues under stress (Kamruzzaman et al. 2025). More recent studies have applied association mapping to traits such as plant height (Kartseva et al. 2024), root system architecture (Siddiqui et al. 2023), nodal root growth angle (Siddiqui et al. 2025), and stomatal traits (Liu et al. 2025). The use of multilocus models has facilitated the identification of genomic regions governing grain micronutrient content (Negi et al. 2025). These findings underscore the utility of GWAS in elucidating the genetic basis of traits critical for wheat improvement.

Complementing these genomic approaches, digital phenotyping now enables high-throughput, non-invasive monitoring of physiological and morphological traits in wheat. Spectral vegetation indices are particularly valuable for assessing drought stress, as they detect early physiological changes often preceding visible morphological symptoms. For instance, NDVI serves as a proxy for canopy greenness and biomass, reflecting yield potential and water-use efficiency, while PRI captures dynamic changes in photosynthetic efficiency and energy dissipation. Similarly, MCARI and SIPI provide insights into pigment composition and chlorophyll degradation. Integrating these spectral indices as digital phenotypes into breeding programs enables rapid identification of stress-resilient wheat genotypes through GWAS. Recent studies have demonstrated that GWAS using spectral vegetation and hyperspectral indices can resolve genomic regions underlying stay-green behavior, moisture-deficit tolerance and grain yield in wheat and durum wheat (Yu et al. 2024; Puttamadanayaka et al. 2020; Mérida-García et al. 2024). Moreover, analyses of synthetic wheat under salinity stress have shown that spectral reflectance indices exhibit substantial genetic variation and heritability and are tightly associated with physiological traits and grain yield, supporting their use as selection proxies in stress-prone environments (Vafadar et al. 2025). Hitherto, indigenous wheat diversity regarding essential vegetative traits has remained largely genetically uncharacterized under the natural rainfed conditions of Azerbaijan. This region is particularly significant as it lies within the center of wheat origin and represents a reservoir of untapped primary genetic resources.

The present study aims to integrate digital phenotyping with GWAS approach to elucidate the genetic architecture controlling spectral vegetation indices in wheat under water-limited conditions. The identified SNP markers and candidate genes can serve as new genetic resources for marker-assisted selection (MAS) in breeding programs targeting the development of drought-tolerant varieties adapted to the dryland environments by indirect selection.

## Material and methods

### Plant material

The plant material utilized in the present study consists of a diversity set of 187 genotypes of bread wheat (*Triticum aestivum* L.) representing a wide geographic origin (Supplementary Table S1). This collection was designed to capture a broad range of phenotypic and genetic diversity relevant for drought adaptation under rainfed conditions.

### Experimental design and growth condition

Field experiments were conducted during the 2022/2023 growing season at the Gobustan Regional Experimental Station of the Research Institute of Crop Husbandry, Azerbaijan (800 m a.s.l.). The soil in this region is classified as light chestnut with a slightly alkaline pH. It is characterized by low humus content (1.5-1.8% in the topsoil), low nitrogen and phosphorus availability, and moderate potassium levels. The hydro-meteorological conditions for the 2022/2023 growing season are presented in detail in Supplementary Table S2. Total precipitation during the cropping season was 356.3 mm, which is below the long-term average of 406.0 mm. A critical moisture deficit occurred in June, with only 4.1 mm of rainfall accompanied by relatively high temperatures, causing pronounced drought stress during the reproductive stages of wheat development. Each experimental plot measured 1.5 × 1.0 m (1.5 m²), with rows spaced 15 cm apart. Sowing was performed at a density of 450 germinating seeds per m². The experiment followed a randomized complete block design with three replications per genotype.

### Evaluation of vegetation indices

Phenological measurements were conducted during the first ten days of June 2023, at the milk ripeness stage. Spectral reflectance of flag leaves was assessed using a portable PolyPen RP400/RP410 spectro-radiometer (Photon Systems Instruments, Czech Republic). This device features an internal xenon incandescent light source (380-1050 nm) and detectors for both UV/VIS and NIR (760-900 nm) regions. Spectral data were normalized against a white calibration standard using the formula:

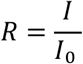

where R is the reflectance, I denotes the signal from the sample, and I_0_ represents the reference signal from the calibration standard. Pseudo-absorbance values were calculated as:

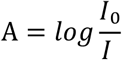

A total of 25 vegetation indices were calculated. Spectral reflectance measurements were conducted with ten biological replicates per genotype. The list of vegetation indices used in this study, including abbreviations, equations and original sources, are provided in Supplementary Table S3.

### Broad sense heritability (*H*^2^)

Descriptive statistics were calculated using R software (v3.6). A mixed linear model (MLM) and two-way ANOVA were performed using the ‘nlme’ and ‘emmeans’ packages to estimate variance components and compare means, considering cultivar and replication as random effects. The variance components were used for calculating broad sense heritability (H^2^). The following formula was used for estimating H^2^:

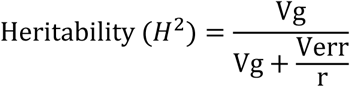

Where, Vg is genotypic variance and Verr is the error variance and r is the number of replications.

### Genotyping of the diversity-set

The wheat accessions were genotyped by TraitGenetics GmbH (Gatersleben, Germany) using the Illumina Infinium XT 25K SNP array. An initial set of 24,146 SNPs was subjected to quality control, which included the removal of monomorphic markers, SNPs with missing data, and markers with a minor allele frequency (MAF) below 5%. This filtering resulted in a final dataset of 19,737 high-quality polymorphic SNPs used for all downstream analyses. Details of these markers are described in Zakieh et al. (2021, 2023).

### GWAS

Genome-wide association mapping was performed following the GRAMMAR method described by Aulchenko et al. (2007). For the analysis, the first three principal components were included as co-factors to control for population structure in the analysis additionally. The methods of the analysis were used and described in detail by Reinert et al. (2016) and Naz et al. (2017). We used a linear mixed model to calculate the QTLs as presented below:

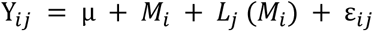

where Y*_ij_* is the phenotypic value; μ is the general mean; *M_i_* is the fixed effect of *i*-th marker genotype; *Lj* (*M_i_*) is the random effect of *j*-th Cultivar nested within *i*-th marker genotype and ε*_ij_* is the residual. Significant markers were detected following a correction using the probability of false discovery rate (FDR), implemented in the SAS procedure PROC MULTTEST according to Benjamini and Yekutieli (2005). To account for multiple testing across all SNPs, genome-wide significance was additionally assessed using a Bonferroni correction based on the total number of markers tested (P ≤ 2.53 × 10⁻⁶).

### Determination of genes colocalizing with QTL

To identify putative candidate genes underlying significant QTL, lead SNP markers were anchored to the wheat reference genome (IWGSC RefSeq v2.1) based on their physical positions in the Ensembl-plant database (Bolser et al. 2017). Candidate gene mining was performed using the BioMart interface (Smedley et al. 2015). A fixed-window approach was applied by defining a ±500 kb genomic interval on both sides of each lead SNP. All annotated gene models whose start or end coordinates fell within this interval were retrieved and considered as putative candidate genes. Functional annotation of candidate genes was obtained based on Gene Ontology (GO) terms describing molecular functions and biological processes, using information available in the UniProt database (The UniProt Consortium 2023).

## Results

### Overall variation of spectral vegetation indices and significant marker trait associations

Broad-sense heritability (*H*^2^) was calculated to estimate the genetic variation and genetic dependence of individual vegetation indices. This analysis revealed a broad spectrum of genetic variation with *H*^2^ estimates ranging from moderate (0.19) to high (0.95) (Fig. 1).

**Fig. 1.**
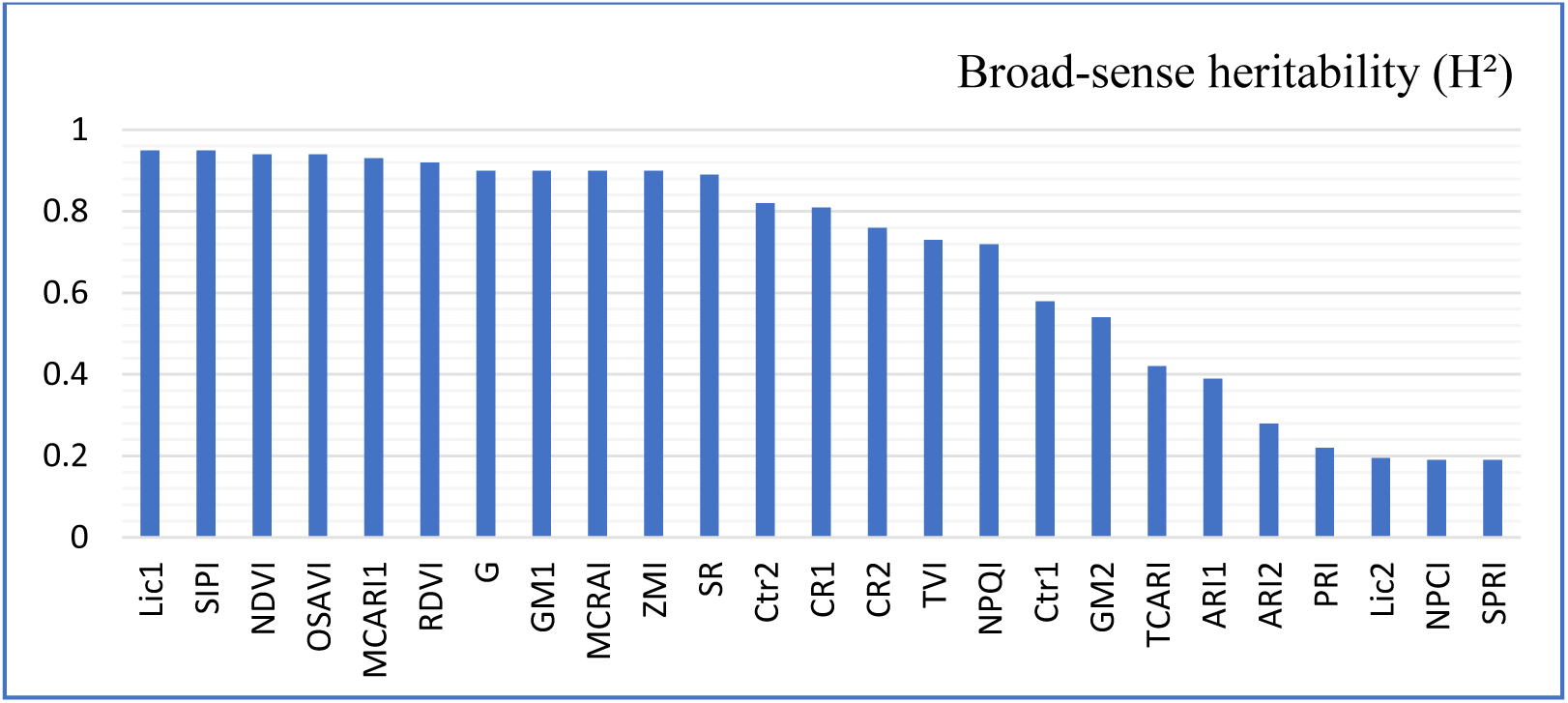
Broad-sense heritability (*H*²) estimates for spectral indices in a diverse wheat panel.

These phenotypic data were compared with 19,737 high-quality SNPs to identify significant marker-trait associations across the genome. These significant marker-trait associations were initially selected based on FDR, which resulted in 4,438 significant SNPs for all selected traits across the genome (Fig. 2). The significant markers were then validated employing Bonferroni correction, which yielded 812 MTAs across the genome. The chromosomal positions of these QTLs are presented in Figure 3, which shows the distribution of QTLs across the 21 wheat chromosomes as well as some hotspots with major effects. Indices related to canopy greenness and structure (e.g., NDVI, OSAVI, Lic1, RDVI, SIPI) exhibited the highest frequency of significant SNPs, whereas pigment-ratio and stress-related indices (e.g., NPCI, SPRI, Lic2, and PRI) resulted in minimal or no significant signals. Among these QTL effects, we established a list of the strongest marker-trait associations which is presented in Table 1.

**Fig. 2.**
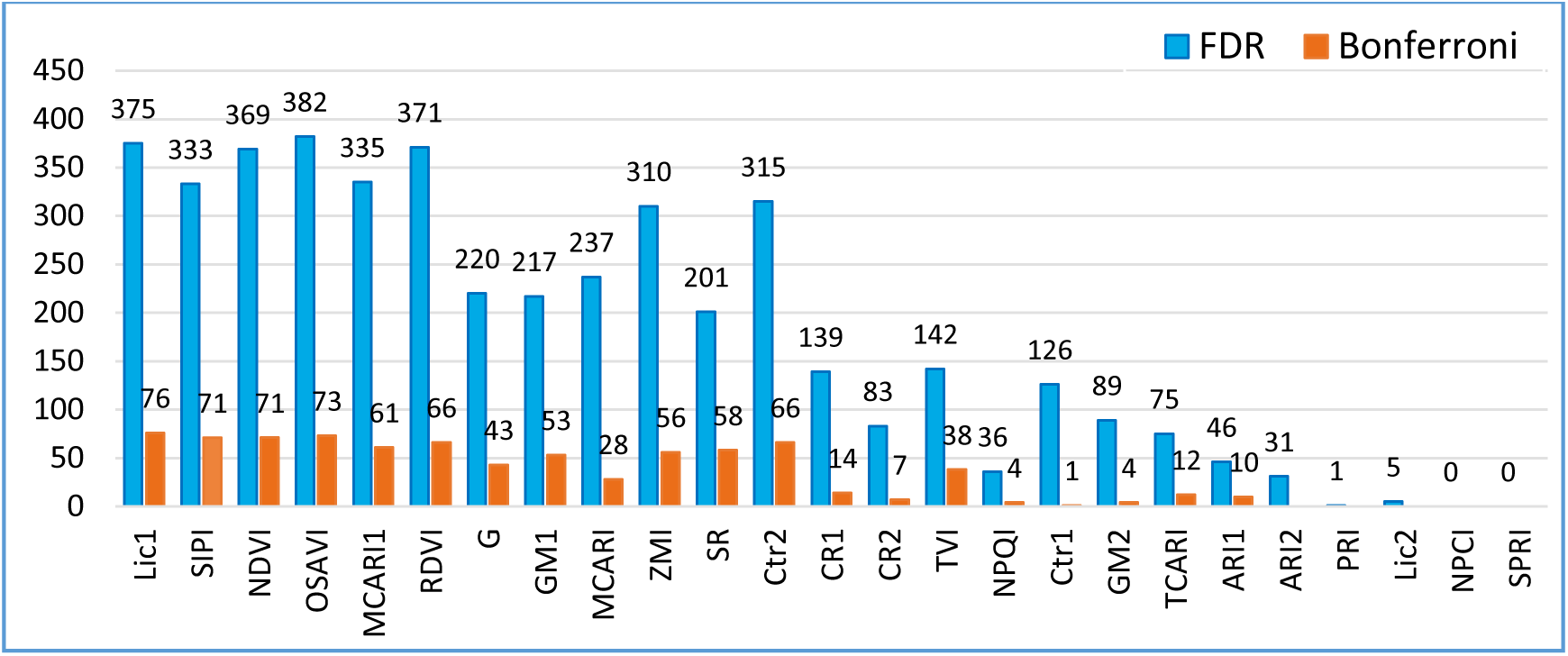
Number of significant MTAs passing the thresholds of FDR and Bonferroni corrections.

**Fig. 3.**
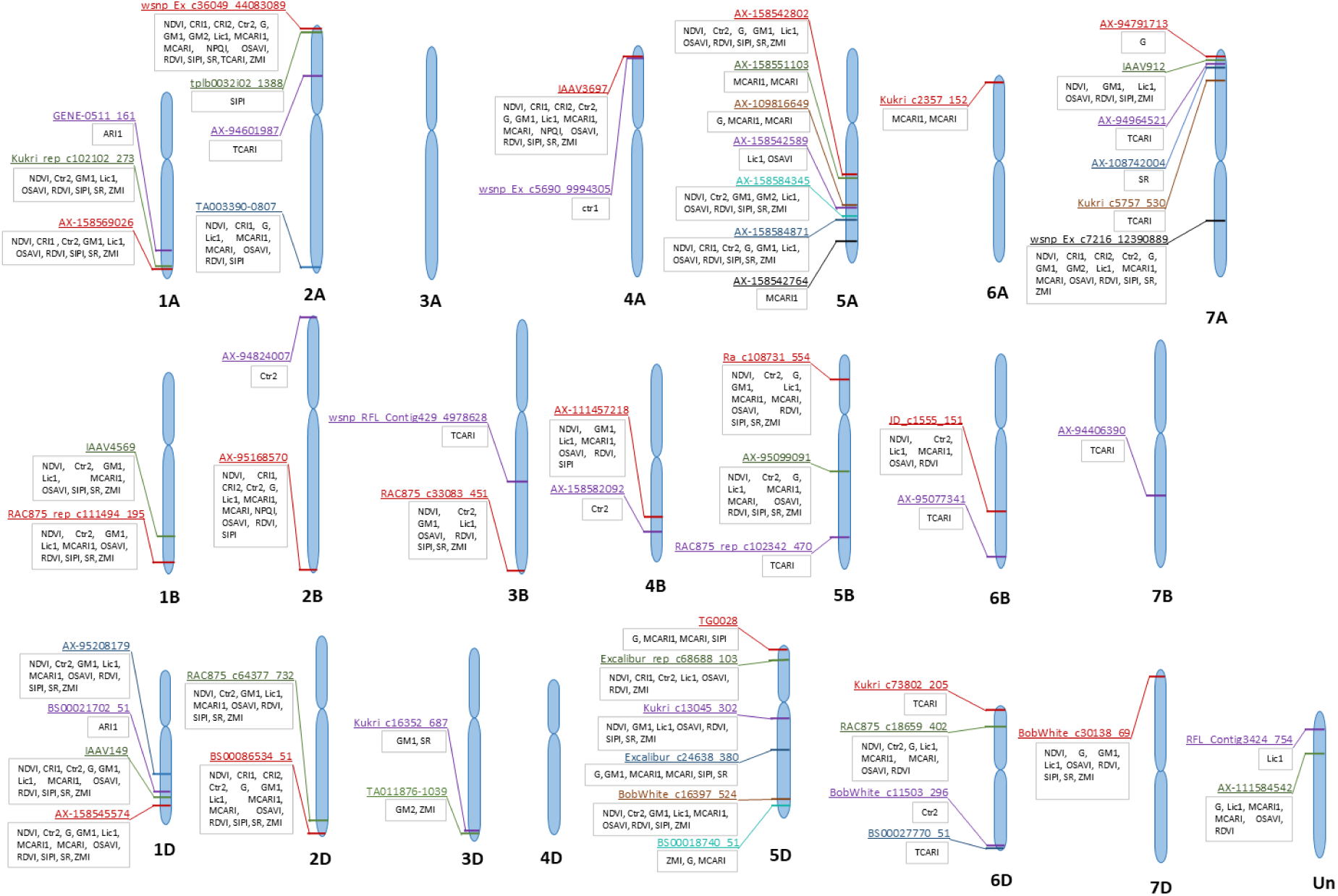
Genome-wide localization of major QTLs for spectral vegetation indices in wheat. Significant MTAs identified at the Bonferroni-corrected threshold are projected onto their physical positions on the 21 wheat chromosomes (1A-7D) and unassigned contigs according to the IWGSC RefSeq v2.1 physical map. Each SNP is annotated with its marker ID and the specific spectral indices it regulates. Colored connectors represent QTL clusters shared across multiple indices, highlighting pleiotropic loci and multi-trait association hotspots.

**Table 1.**
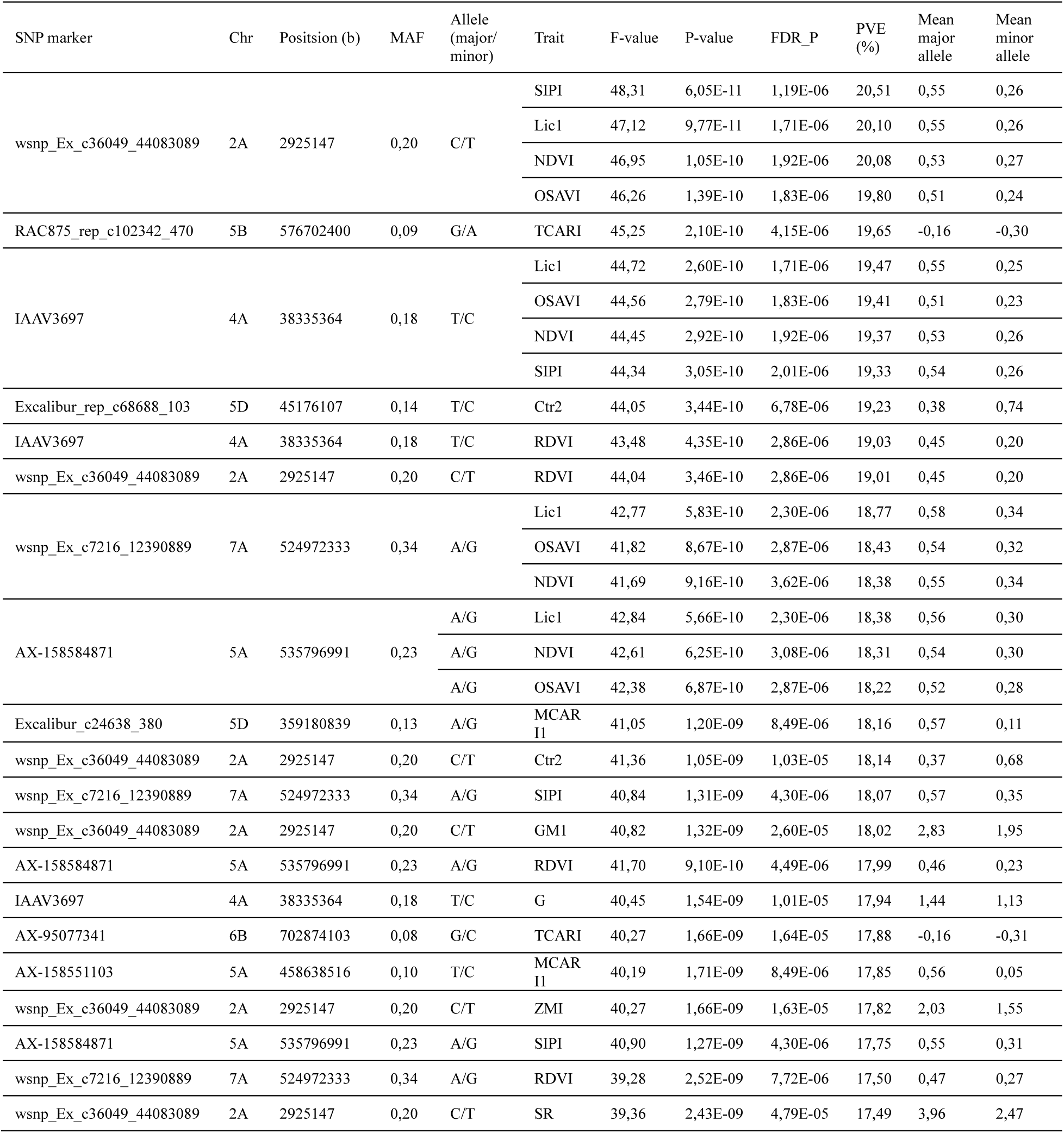
Top 20 most significant QTL for vegetation indices in wheat.

The description of significant MTAs for individual vegetation indices are described below:

### NDVI

As the most widely used greenness metric, NDVI exhibited the highest heritability in our dataset (*H*² = 0.94). NDVI also revealed the one of the highest and strongest significant MTAs. In total, 71 SNPs exceeded the Bonferroni threshold, spanning 16 chromosomes and explaining 5.27–20.08% of the genotypic variance (Supplementary Table S4). To summarize the genome-wide signal, we applied LD-based clumping to all SNPs showing statistically significant association, which resolved them into 76 independent peak loci, 28 of which remained above the Bonferroni threshold (Fig. 4a, b). The largest-effect locus was on chromosome 2A at wsnp_Ex_c36049_44083089, where the major allele (C) increased NDVI values relative to the minor allele (T), with mean values of 0.53 vs 0.27 (Table 1). Additional major-effect loci were identified on chromosomes 4A (IAAV3697), 5A (AX-158584871), 5D (Excalibur_rep_c68688_103) and 7A (wsnp_Ex_c7216_12390889), representing the strongest associations after the lead SNP and ranking among the top five in terms of explained variance (PVE ≥ 16.4%). Across the 76 peak loci, the major allele was favorable at 59 loci, whereas the minor allele increased the trait at 17 loci (Supplementary Table S5). Overall, the significant SNP markers were distributed across A, B and D genomes (Fig. 4c).

**Fig. 4.**
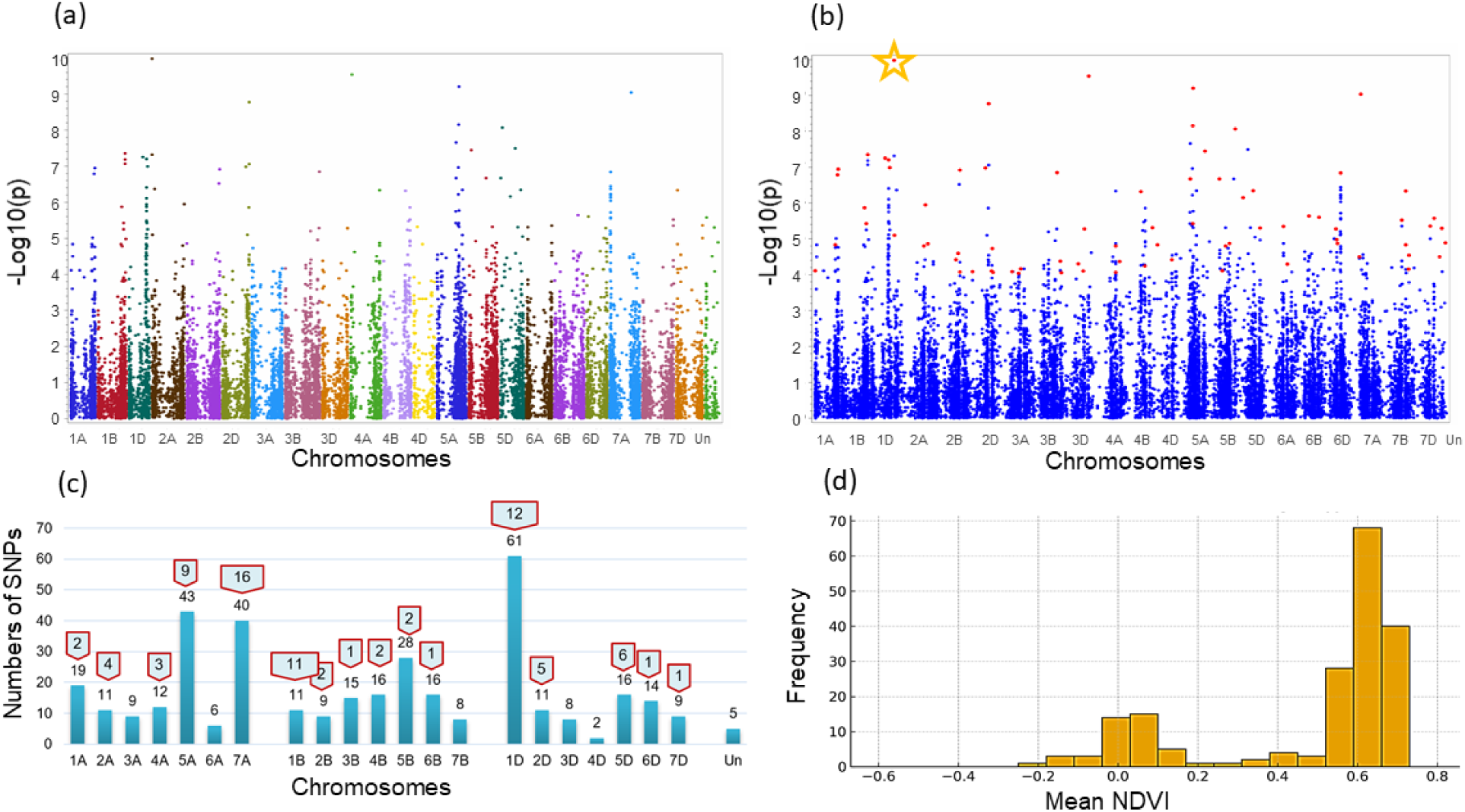
Significant genome-wide associations for NDVI. (a) Manhattan plot of SNP-based GWAS for NDVI across the wheat genome. (b) Manhattan plot of LD-clumped NDVI associations, highlighting LD-independent peak markers (red) selected based on p-value and local LD structure; the star marks the largest-effect locus on chromosome 2A at wsnp_Ex_c36049_44083089. (c) Chromosomal distribution of significant NDVI-associated SNPs; blue bars show the total number of SNPs with statistically significant association per chromosome (FDR), and red labels indicate the subset of Bonferroni-significant SNPs. (d) Distribution of NDVI phenotypic values across the wheat panel; frequency denotes the number of genotypes.

### ARI1 and ARI2

ARI1, reflecting anthocyanin accumulation, displayed moderate broad-sense heritability (*H*² = 0.39). Ten Bonferroni-significant SNPs were detected on chromosomes 1A and 1D, each explaining 3.24–12.67% of the variance (Supplementary Table S4). LD-based clumping of all SNPs showing statistically significant association grouped these into six independent peak loci, two of which exceeded the Bonferroni threshold. The top association was on chromosome 1A at GENE-0511_161, where the major allele (A) increased the trait relative to the minor allele (G), with mean values of −0.06 vs −0.20. A second major-effect locus on chromosome 1D at BS00021702_51 had PVE = 12.53% (Supplementary Table S5). At all six peak loci, the major allele was associated with higher values.

The secondary anthocyanin-related index, ARI2, exhibited low-to-moderate heritability (*H*² = 0.28). No SNP reached the Bonferroni threshold, but 25 SNPs on eight chromosomes were significant at FDR q ≤ 0.05, each explaining 8.76–10.18% of the phenotypic variance. LD-based clumping of all SNPs showing statistically significant association defined eight independent peak loci. The strongest signals were on chromosome 1D at BS00021702_51 and on chromosome 1A at CAP11_c3464_68, each explaining approximately 10.18% of the variance. Across all FDR-significant loci, the major allele was consistently associated with higher ARI2 values than the minor allele (Supplementary Table S4).

### CRI1 and CRI2

The carotenoid-related reflectance index CRI1 was highly heritable (*H*² = 0.81). Fourteen SNPs on nine chromosomes exceeded the Bonferroni threshold, with PVE values ranging from 11.35% to 15.80% (Supplementary Table S4). LD-based clumping of all SNPs showing statistically significant association revealed 38 independent peak loci, including ten Bonferroni-significant peaks (Supplementary Table S5). The strongest association was on chromosome 4A at IAAV3697, where the major allele (T) increased values relative to the minor allele (C), with mean values of 1.99 vs 0.90. Additional major-effect loci occurred on chromosomes 2A (wsnp_Ex_c36049_44083089), 7A (wsnp_Ex_c7216_12390889), 2B (AX-95168570) and 1A (AX-158569026), representing the strongest associations after the lead SNP and ranking among the top five loci for explained variance (PVE ≥ 12.25%). Across the 38 peak loci, the major allele increased the trait at 33 loci and the minor allele at five.

CRI2 also showed a substantial genetic component (*H*² = 0.76). Seven SNPs on five chromosomes exceeded the Bonferroni threshold, explaining 10.24–15.09% of the variance (Supplementary Table S4). LD-based clumping of all SNPs showing statistically significant association defined 13 independent peak loci, five of which were Bonferroni-significant (Supplementary Table S5). The strongest signal was again on chromosome 4A at IAAV3697, where the major allele (T) increased values compared with the minor allele (C), with mean values of 1.85 vs 0.81. Additional major-effect loci were detected on chromosomes 2A (wsnp_Ex_c36049_44083089), 7A (wsnp_Ex_c7216_12390889), 2B (AX-95168570) and 2D (BS00086534_51), ranking among the top five loci for explained variance (PVE ≥ 10.24%). Across the 13 peak loci, the major allele was favorable at 12 loci and the minor allele at one.

### Ctr1 and Ctr2

Ctr1 exhibited moderate heritability (*H*² = 0.58). One SNP on chromosome 4A exceeded the Bonferroni threshold, explaining 11.67% of the variance (Supplementary Table S4). LD-based clumping of all SNPs showing statistically significant association identified three independent peak loci. The leading association on 4A at wsnp_Ex_c5690_9994305 showed higher values for the major allele (G) than for the minor allele (A), with mean values of 1.28 vs 1.13. Additional loci on chromosomes 2D (Excalibur_c42413_442) and 5D (TG0028) further contributed to the observed variation. At all three peak loci, the major allele increased the index.

In contrast, Ctr2 was highly heritable (*H*² = 0.82). Sixty-six SNPs surpassed the Bonferroni threshold across 15 chromosomes, explaining 4.76–19.23% of the variance (Supplementary Table S4). LD-based clumping of all SNPs showing statistically significant association grouped these signals into 68 independent peak loci, including 26 Bonferroni-significant peaks (Supplementary Table S5). The strongest locus on chromosome 5D at Excalibur_rep_c68688_103 showed lower values for the major allele (T) than for the minor allele (C), with mean values of 0.38 vs 0.74. Additional major-effect loci were detected on chromosomes 2A (wsnp_Ex_c36049_44083089), 5A (AX-158584871), 7A (wsnp_Ex_c7216_12390889) and 4A (IAAV3697), representing the strongest associations after the lead SNP (PVE ≥ 16.41%). Across the 68 peak loci, the major allele increased the trait at only 11 loci, while the minor allele was favorable at 57.

### G index

The G index showed high heritability (*H*² = 0.90). Forty-three SNPs exceeded the Bonferroni threshold, with 42 mapping to 11 chromosomes and one to an unassigned contig, explaining 5.57–17.94% of the variance (Supplementary Table S4). LD-based clumping of all SNPs showing statistically significant association yielded 64 independent peak loci, including 20 Bonferroni-significant peaks (Supplementary Table S5). The strongest association was on chromosome 4A at IAAV3697, where the major allele (T) increased values relative to the minor allele (C), with mean values of 1.44 vs 1.13. Additional major-effect loci were detected on chromosomes 7A (wsnp_Ex_c7216_12390889), 2A (wsnp_Ex_c36049_44083089), 2D (BS00086534_51) and 5D (Excalibur_c24638_380), representing the strongest associations after the lead SNP and ranking among the top five loci for explained variance. Across the 64 peak loci, the major allele increased the trait at 48 loci and the minor allele at 16.

### GM1 and GM2

GM1 was highly heritable (*H*² = 0.90). Fifty-three Bonferroni-significant SNPs were identified on 14 chromosomes, explaining 6.83–18.02% of the variance (Supplementary Table S4). LD-based clumping of all SNPs showing statistically significant association identified 61 independent peak loci, including 24 Bonferroni-significant peaks (Supplementary Table S5). The strongest locus was on chromosome 2A at wsnp_Ex_c36049_44083089, where the major allele (C) increased values relative to the minor allele (T), with mean values of 2.83 vs 1.95. Additional major-effect loci were located on chromosomes 7A (wsnp_Ex_c7216_12390889), 4A (IAAV3697), 5A (AX-158584871) and 1D (AX-95208179), ranking among the top five loci for explained variance (PVE ≥ 13.88%). Across the 61 peak loci, the major allele increased the index at 49 loci and the minor allele at 12.

GM2 showed moderate heritability (*H*² = 0.54). Four Bonferroni-significant SNPs on four chromosomes explained 7.83–12.89% of the variance (Supplementary Table S4). LD-based clumping of all SNPs showing statistically significant association yielded three independent peak loci, all genome-wide significant. The strongest association was on chromosome 7A at wsnp_Ex_c7216_12390889, where the major allele (A) increased values relative to the minor allele (G), with mean values of 3.04 vs 2.25. Additional loci on chromosomes 2A (wsnp_Ex_c36049_44083089) and 5A (AX-158584345) further contributed to the observed variation. Among the three peak loci, the major allele was favorable at two and the minor allele at one.

### Lic1 and Lic2

Lic1 showed very high heritability (*H*² = 0.95). Seventy-six SNPs exceeded the Bonferroni threshold, spanning 16 chromosomes and one unassigned scaffold and explaining 5.42–20.10% of the variance (Supplementary Table S4). LD-based clumping of all SNPs showing statistically significant association identified 74 independent peak loci, including 31 Bonferroni-significant peaks (Supplementary Table S5). The strongest association was on chromosome 2A at wsnp_Ex_c36049_44083089, where the major allele (C) increased values compared with the minor allele (T), with mean values of 0.55 vs 0.26. Additional major-effect loci occurred on chromosomes 4A (IAAV3697 and wsnp_Ex_rep_c67145_65628860), 5A (AX-158584871), 5D (Excalibur_rep_c68688_103) and 7A (wsnp_Ex_c7216_12390889), representing the strongest associations after the lead SNP (PVE ≥ 16.73%). Across the 74 peak loci, the major allele increased the trait at 57 loci and the minor allele at 17.

Lic2 had low broad-sense heritability (*H*² = 0.20) and did not show any significant genetic associations at the applied thresholds.

### MCARI and MCARI1

MCARI displayed high heritability (*H*² = 0.90). Twenty-eight SNPs exceeded the Bonferroni threshold, spanning 11 chromosomes and one unassigned scaffold and explaining 5.57–16.77% of the variance (Supplementary Table S4). LD-based clumping of all SNPs showing statistically significant association identified 53 independent peak loci, including 17 Bonferroni-significant peaks (Supplementary Table S5). The leading locus was on chromosome 2D at Excalibur_c42413_442, where the major allele (T) increased values relative to the minor allele (C), with mean values of 0.072 vs 0.003. Additional major-effect loci were detected on chromosomes 4A (IAAV3697), 5D (Excalibur_c24638_380), 5A (AX-158551103) and 2B (AX-95168570), ranking among the top five loci for explained variance (PVE ≥ 15.48%). Across the 53 peak loci, the major allele increased the index at 47 loci and the minor allele at six.

MCARI1 was also highly heritable (*H*² = 0.93). Sixty-one Bonferroni-significant SNPs on 14 chromosomes and one unassigned scaffold explained 4.94–18.16% of the variance (Supplementary Table S4). LD-based clumping of all SNPs showing statistically significant association revealed 74 independent peak loci, including 26 Bonferroni-significant peaks (Supplementary Table S5). The strongest association was on chromosome 5D at Excalibur_c24638_380, where the major allele (A) increased values relative to the minor allele (G), with mean values of 0.57 vs 0.11. Additional major-effect loci were located on chromosomes 5A (AX-158551103), 4A (IAAV3697), 2A (wsnp_Ex_c36049_44083089) and 7A (wsnp_Ex_c7216_12390889) representing the strongest associations after the lead SNP (PVE ≥ 16.78%). Across the 74 peak loci, the major allele increased the trait at 60 loci and the minor allele at 14.

### NPQI

NPQI showed moderately high heritability (*H*² = 0.72). Four SNPs exceeded the Bonferroni threshold on chromosomes 2A, 2B and 4A, explaining 11.23–12.85% of the variance (Supplementary Table S4). LD-based clumping of all SNPs showing statistically significant association grouped these into three independent peak loci, all genome-wide significant (Supplementary Table S5). The strongest association was on chromosome 4A at IAAV3697, where the minor allele (C) increased values relative to the major allele (T), with mean values of −0.011vs −0.031. Additional loci on chromosomes 2A (wsnp_Ex_c36049_44083089) and 2B (AX-95168570) also contributed to the observed variation. At all three peak loci, the minor allele increased the index.

### NPCI

NPCI exhibited low heritability (*H*² = 0.19) and no significant associations.

### OSAVI

OSAVI was very highly heritable (*H*² = 0.94). Seventy-three SNPs exceeded the Bonferroni threshold, mapping to 16 chromosomes and one unassigned scaffold and explaining 5.43–19.80% of the variance (Supplementary Table S4). LD-based clumping of all SNPs showing statistically significant association identified 73 independent peak loci, including 30 Bonferroni-significant peaks (Supplementary Table S5). The strongest association was on chromosome 2A at wsnp_Ex_c36049_44083089, where the major allele (C) increased values relative to the minor allele (T), with mean values of 0.51 vs 0.24. Additional major-effect loci occurred on chromosomes 4A (IAAV3697), 5A (AX-158584871), 5D (Excalibur_rep_c68688_103) and 7A (wsnp_Ex_c7216_12390889), representing the strongest associations after the lead SNP (PVE ≥ 16.85%). Across the 73 peak loci, the major allele increased the trait at 59 loci and the minor allele at 14.

### PRI

PRI had low broad-sense heritability (*H*² = 0.22) and no significant associations.

### RDVI

RDVI exhibited very high heritability (*H*² = 0.92). Sixty-six SNPs exceeded the Bonferroni threshold, spanning 16 chromosomes and one unassigned scaffold and explaining 5.40–19.03% of the variance (Supplementary Table S4). LD-based clumping of all SNPs showing statistically significant association identified 69 independent peak loci, including 28 Bonferroni-significant peaks (Supplementary Table S5). The strongest association was on chromosome 4A at IAAV3697, where the major allele (T) increased values relative to the minor allele (C), with mean values of 0.45 vs 0.20. Additional major-effect loci were detected on chromosomes 2A (wsnp_Ex_c36049_44083089), 5A (AX-158584871), 5D (Excalibur_rep_c68688_103) and 7A (wsnp_Ex_c7216_12390889), representing the strongest associations after the lead SNP and ranking among the top five loci for explained variance (PVE ≥ 17.44%). Across the 69 peak loci, the major allele increased the trait at 55 loci and the minor allele at 14.

### SIPI

SIPI showed very high heritability (*H*² = 0.95). Seventy-one Bonferroni-significant SNPs on 14 chromosomes explained 5.13–20.51% of the variance (Supplementary Table S4). LD-based clumping of all SNPs showing statistically significant association yielded 75 independent peak loci, including 28 Bonferroni-significant peaks (Supplementary Table S5). The strongest association was on chromosome 2A at wsnp_Ex_c36049_44083089, where the major allele (C) increased values relative to the minor allele (T), with mean values of 0.55 vs 0.26. Additional major-effect loci were located on chromosomes 4A (IAAV3697), 5A (AX-158584871), 5D (Excalibur_c24638_380) and 7A (wsnp_Ex_c7216_12390889), ranking among the top five loci (PVE ≥ 15.27%). Across the 75 peak loci, the major allele increased the trait at 59 loci and the minor allele at 16.

### SPRI

SPRI had low heritability (*H*² = 0.19) and no significant associations.

### SR

SR showed high heritability (*H*² = 0.89). Fifty-eight SNPs exceeded the Bonferroni threshold across 13 chromosomes, explaining 6.10–17.49% of the variance (Supplementary Table S4). LD-based clumping of all SNPs showing statistically significant association revealed 56 independent peak loci, including 23 Bonferroni-significant peaks (Supplementary Table S5). The strongest association was on chromosome 2A at wsnp_Ex_c36049_44083089, where the major allele (C) increased values relative to the minor allele (T), with mean values of 3.96 vs 2.47. Additional major-effect loci were detected on chromosomes 7A (wsnp_Ex_c7216_12390889), 4A (IAAV3697), 1D (IAAV149) and 5A (AX-158584871), representing the strongest associations after the lead SNP (PVE ≥ 13.69%). Across the 56 peak loci, the major allele increased the trait at 45 loci and the minor allele at 11.

### TCARI

TCARI showed moderate heritability (*H*² = 0.42). Twelve SNPs on seven chromosomes surpassed the Bonferroni threshold and explained 10.18–19.65% of the phenotypic variance (Supplementary Table S4). LD-based clumping of all SNPs showing statistically significant association identified 24 independent peak loci, including nine Bonferroni-significant peaks (Supplementary Table S5). The strongest association was on chromosome 5B at RAC875_rep_c102342_470, where the major allele (G) increased values relative to the minor allele (A), with mean values of −0.16 vs −0.30. Additional major-effect loci were located on chromosomes 6B (AX-95077341), 7B (AX-94406390), 2A (AX-94601987), ranking among the top five loci (PVE ≥ 16.70%). Among the 24 peak loci, the major allele increased the index at 23 loci and the minor allele at one.

### TVI

TVI had intermediate heritability (*H*² = 0.73). Thirty-eight SNPs exceeded the Bonferroni threshold across 12 chromosomes and one unassigned scaffold and explained 4.28–19.48% of the variance (Supplementary Table S4). LD-based clumping of all SNPs showing statistically significant association identified 34 independent peak loci, including 18 Bonferroni-significant peaks (Supplementary Table S5). The strongest association was on chromosome 5D at Excalibur_c24638_380, where the major allele (A) increased values relative to the minor allele (G), with mean values of 2.06 vs −0.41. Additional major-effect loci were detected on chromosomes 5A (AX-158551103), 5B (AX-95099091), 2A (AX-94406025) and 6D (BobWhite_c11503_296), representing the strongest associations after the lead SNP (PVE ≥ 14.04%). Across the 34 peak loci, the major allele increased the trait at 32 loci and the minor allele at two.

### ZMI

ZMI was highly heritable (*H*² = 0.90). A total of 56 SNPs exceeded the Bonferroni threshold, spanning 13 chromosomes and explaining 7.91–17.82% of the variance (Supplementary Table S4). LD-based clumping of all SNPs showing statistically significant association revealed 72 independent peak loci, including 24 Bonferroni-significant peaks (Supplementary Table S4). The strongest association was on chromosome 2A at wsnp_Ex_c36049_44083089, where the major allele (C) increased values relative to the minor allele (T), with mean values of 2.03 vs 1.55. Additional major-effect loci were located on chromosomes 7A (wsnp_Ex_c7216_12390889), 4A (IAAV3697), 1B (RAC875_rep_c111494_195) and 1D (AX-95208179), ranking among the top five loci for explained variance (PVE ≥ 14.91%). Across the 72 peak loci, the major allele increased the trait at 55 loci and the minor allele at 17.

### Candidate gene localization at the 2A QTL region

Within the ±500 kb physical interval centered on the lead SNP *wsnp_Ex_c36049_44083089* (2A: 2,925,147 bp), the marker is situated approximately 4.2 kb downstream of *TraesCS2A02G006800.1* and 39 kb upstream of *TraesCS2A02G006900.1*. Both genes encode NB-ARC proteins containing leucine-rich repeat (LRR) and disease-resistance domains (Supplementary Table S6). In total, this genomic interval harbors five NB-ARC/LRR-type genes (*TraesCS2A02G004900.1*, *G006500.1*, *G006800.1*, *G006900.1*, *G007000.1*). Notable stress-related candidates identified in this region include *TraesCS2A02G005500.1*, which encodes a LEA_2/NDR1-like protein, and *TraesCS2A02G006100.1*, encoding a lectin receptor-like serine/threonine kinase featuring S-domain, bulb-type lectin, and PAN/Apple domains. Furthermore, the locus is enriched with genes involved in specialized and primary metabolism, including a cluster of cytochrome P450 monooxygenases (*TraesCS2A02G005100.1*, *G005800.1*, *G005900.1*, *G006300.1*, *G006400.1*), a terpene synthase (*TraesCS2A02G006000.1*), secondary-metabolite acyltransferases (*TraesCS2A02G006200.1*, *G006700.1*), a hexokinase-like phosphotransferase (*TraesCS2A02G005200.1*), and proton-dependent oligopeptide/MFS transporters (*TraesCS2A02G007100.1*, *G007500.1*).

## Discussion

Enhancing drought tolerance in wheat is a critical breeding objective for rainfed agriculture, which necessitates the identification of robust genetic markers and a deeper understanding of the underlying physiological mechanisms. This study integrated high-throughput digital phenotyping with whole-genome SNP data of a unique wheat diversity panel evolved and selected primarily in the rainfed conditions of Azerbaijan. Azerbaijan belongs to the wheat center of diversity which putatively harbors untapped genetic resources relevant for rainfed wheat production. Previous work on Azerbaijani bread wheat has documented valuable drought-adaptive variation, spanning biochemical and cellular responses to water deficit as well as informative non-invasive vegetation indices and drought-related gene expression (Allahverdiyev et al. 2025; Rustamova et al. 2021; Aliyeva et al. 2024). Therefore, we hypothesize that the genome-wide association analysis of primary vegetation indices will deliver new genomic loci and markers relevant for drought stress adaptation for rainfed wheat production worldwide. Our results reveal several key findings: (1) a wide range of broad-sense heritability among vegetation indices, from very high (*H*^2^ > 0.90) to very low (*H*^2^ ∼ 0.19), highlighting the substantial genetic variation of the physiological traits; (2) the identification of numerous significant MTAs, with several major QTL effects, which also exhibit pleiotropic effects for multiple indices.

### Genetic architecture of spectral indices

The wide range of broad-sense heritability estimates observed for the 25 spectral indices indicates that digital phenotypes differ markedly in the strength of their genetic control under drought. Greenness- and structure-related indices such as NDVI, Lic1, OSAVI, GM1, RDVI, SIPI, SR and ZMI showed very high heritability (*H*² up to 0.95), whereas NPCI, PRI, SPRI and Lic2 exhibited low heritability (*H*² ≈ 0.19-0.22). Similar patterns have been reported in other genetic studies, where NDVI and other greenness- and structure-related indices demonstrated high heritability and strong associations with grain yield under drought and heat stress, confirming their utility as stable phenotypic proxies for selection. Whereas indices related to light-use efficiency and pigments showed lower heritability and greater environmental dependency (Babar et al. 2007; Liu et al. 2019; Mérida-García et al. 2024). The high heritability of most greenness indices in our study confirms that they can be used as robust digital traits to capture genetic differences in canopy maintenance and “stay-green” behavior under water-limited conditions.

### A pleiotropic hotspot on chromosome 2A

The most striking result was the identification of a strong pleiotropic locus on chromosome 2A, tagged by wsnp_Ex_c36049_44083089 (Tabe 1). This SNP was among the top associations for 17 vegetation indices, with the proportion of genotypic variance explained ranging from 12.30% to 20.51%. The largest effects were observed for SIPI, Lic1, NDVI and OSAVI, followed by RDVI, GM1, ZMI and SR. At this locus, the major SNP allele (C) increased the values of the key greenness and structural indices relative to the minor allele (T). For a few stress-sensitive ratios, such as Ctr2 and NPQI, the major allele was associated with lower index values, consistent with their inverse scaling. Since NPQI increases during chlorophyll degradation and Ctr2 is sensitive to leaf water stress and structural breakdown, the association of the favorable allele with their lower values directly indicates reduced stress and less senescence in these genotypes (Joynson et al. 2021). The prominence of the 2A hotspot in our panel is consistent with numerous reports highlighting chromosome 2A as a key region for stress adaptation and canopy traits in wheat. Puttamadanayaka et al. (2020) mapped 15 QTLs for NDVI across the genome, including QNdvi2.iari_2A using a backcross inbred population evaluated under moisture-deficit conditions, which explained the highest proportion of phenotypic variance for NDVI. A meta-analysis by Acuña-Galindo et al. (2015) identified a meta-QTL (MQTL11-13) on chromosome 2A that integrates QTL for grain yield under drought and heat, further emphasizing the importance of this region for combined stress tolerance. Consistently, Singh et al. (2025) mapped three stable QTL on chromosome 2A (QHgfr.iiwbr-2A, QHgns.iiwbr-2A, QLgns.iiwbr-2A) under normal and late-sown heat stress, with loci explaining 6.99–12.22% of the phenotypic variance. Taken together, these results suggest that the 2A region harbors genes with major QTL effects which may reveal pleiotropic effects on canopy greenness, pigment maintenance and yield-related physiology under water-limited and heat-prone field conditions.

### Additional multi-trait loci on chromosomes 4A, 5A and 7A

Besides the 2A hotspot, several genomic regions on chromosomes 4A, 5A and 7A contributed strongly to multiple indices (Fig 3). Marker IAAV3697 on 4A showed leading or near-leading effects for Lic1, OSAVI, NDVI, SIPI, RDVI, and G, with PVE values ranging from 17.9% to 19.5%, and consistently higher index values for the major allele. On chromosome 5A, AX-158584871 was among the top QTL for Lic1, NDVI, OSAVI, RDVI and SIPI with PVE values ranging from 17.8% to 18.4%. On 7A, wsnp_Ex_c7216_12390889 repeatedly emerged as a major locus for Lic1, OSAVI, NDVI, SIPI, and RDVI with PVE values 17.5% - 18.8% (Table 1). The multi-trait loci detected on these chromosomes coincide with chromosomal regions previously shown to harbor QTL clusters for drought and heat adaptation. The multi-trait loci we detected fall on chromosomes that have been repeatedly highlighted as hotspots for drought-stress adaptation. In an association panel grown under contrasting Mediterranean rainfed and humid irrigated conditions, Ballesta et al. (2019) identified chromosome 4A as the main QTL-rich region, harbouring 26 SNP–trait associations for drought tolerance indices of grain yield and its components, particularly thousand-kernel weight and kernels per spike. In a bread wheat backcross population, Puttamadanayaka et al. (2020) showed that chromosomes 2A, 5A, 5D, 6A and 6D commonly harboured QTL for moisture-deficit stress tolerance, and mapped 15 NDVI QTL on 2A, 5A, 5D, 6A, 6B, 6D, 2D, 7A and 7B, with seven NDVI QTL sharing confidence intervals with QTL for harvest index, grain yield, biomass, relative water content, days to heading, canopy temperature and SPAD, and QNdvi3.iari_5A co-localizing with a QTL for chlorophyll content. Together, these studies and our results indicate that the 2A, 4A, 5A, 7A chromosome segments represent conserved genomic regions where alleles controlling canopy reflectance, stay-green traits and yield-related performance tend to co-aggregate under water-limited conditions.

### Candidate genes underlying the 2A QTL

The gene composition of the 2A region surrounding wsnp_Ex_c36049_44083089 is consistent with a QTL that integrates immune-related signalling with abiotic stress adaptation under water deficit. In this interval, the lead SNP lies ∼4.2 kb downstream of the NB-ARC–LRR gene *TraesCS2A02G006800.1*, within a tight cluster of NLR-like genes comprising *TraesCS2A02G004900.1, TraesCS2A02G006500.1, TraesCS2A02G006800.1, TraesCS2A02G006900.1* and *TraesCS2A02G007000.1* (Fig. 5), all carrying canonical NB-ARC and LRR signatures together with N-terminal disease-resistance and apoptotic protease–activating factor–like helical domains (Supplementary Table S6). There is now clear evidence that some NLRs can directly enhance abiotic stress tolerance in addition to their canonical roles in disease resistance. For example, the rice NLR PibH8, originally characterized as a blast-resistance homologue of Pib, was recently shown to confer drought resistance by interacting with phenylalanine ammonia-lyase (OsPAL1), protecting it from E3-ligase–mediated degradation, increasing lignin and flavonoid accumulation, and thereby improving drought performance while maintaining blast resistance (Xiang et al. 2025). Likewise, the rice NLR RGA4L was identified as the causal gene of a major chilling-tolerance QTL, where overexpression enhances cold tolerance at both vegetative and reproductive stages and the protein physically interacts with OsHSP90 and the LEA protein OsLEA5, linking an NLR complex to low-temperature signaling (Gan et al. 2025). Genome-wide and evolutionary analyses further indicate that many NLRs are transcriptionally responsive to abiotic stimuli such as drought and temperature stress (Yang et al. 2021), and that disease-resistance–associated miRNAs can be dynamically regulated by heat to modulate NB-LRR transcript abundance (Zang et al. 2019). Together, these studies support the view that the NLR cluster flanking wsnp_Ex_c36049_44083089 is compatible with dual roles in biotic and abiotic stress responses.

**Fig. 5.**
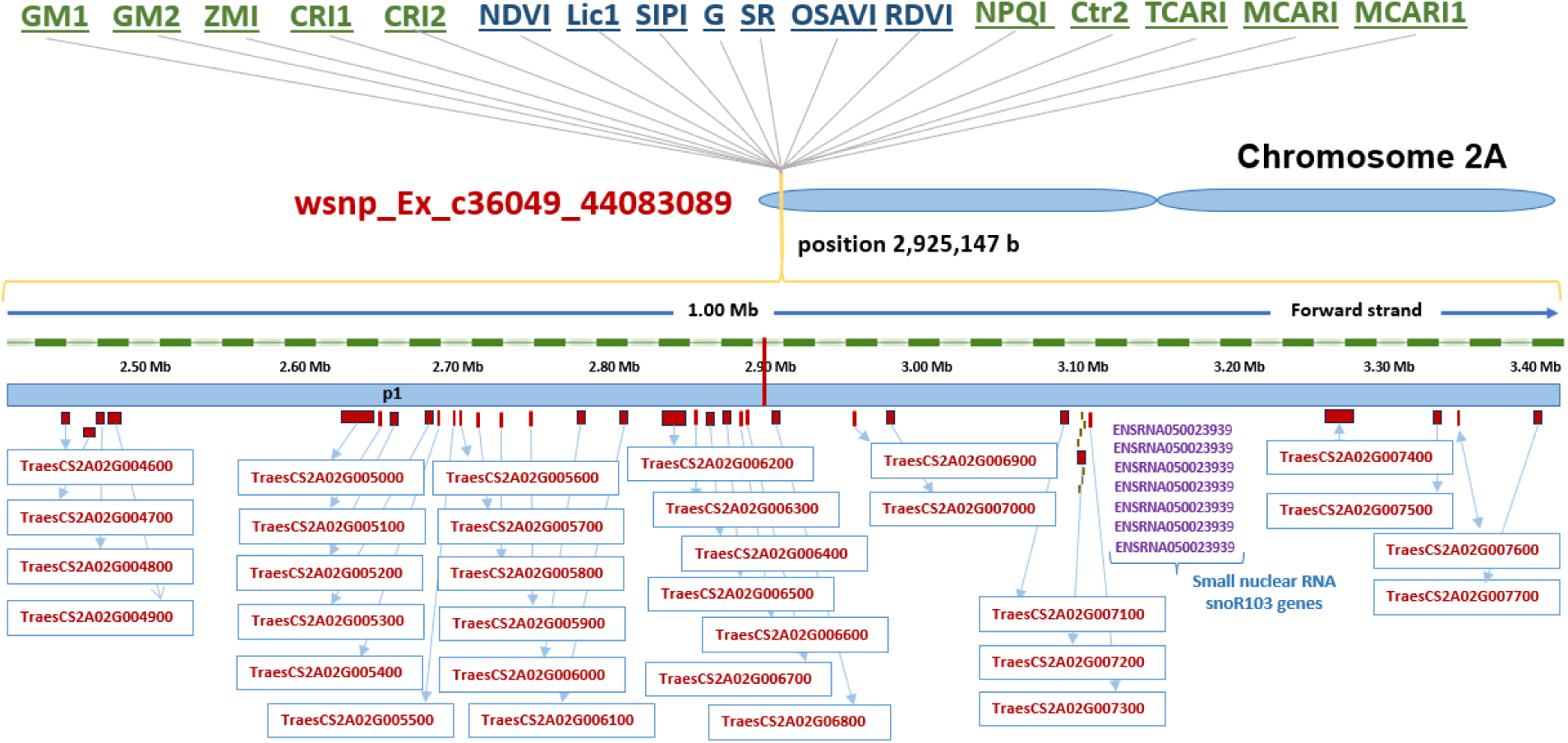
Chromosome 2A QTL region around SNP marker wsnp_Ex_c36049_44083089 and putative candidate genes.

The co-localization of this NLR cluster with a LEA_2/NDR1-like gene in the same interval suggests an additional layer of cross-talk between immune and abiotic stress pathways (Amato et al. 2025). In Arabidopsis, the immune regulator NDR1 has been shown to be required for ABA-dependent drought responses and for proper regulation of plasma-membrane H⁺-ATPases and stomatal behavior, providing a mechanistic link between drought tolerance and defence signalling. Members of the NDR1/HIN1-like (NHL) family carrying LEA_2 motifs are broadly associated with abiotic stress: NHL6, for example, is strongly induced by ABA, osmotic and drought stress, and its overexpression alters ABA sensitivity and abiotic-stress responses, while recent work on NHL12 identified it as a drought-responsive regulator of plasmodesmata permeability and ABA signaling (Bao et al. 2016). These findings make a LEA_2/NDR1-like candidate in the 2A interval a plausible component of signalling modules that integrate water-deficit and defense cues.

The presence of a lectin receptor-like serine/threonine kinase in the same region (*TraesCS2A02G006100.1*) adds a plausible cell-surface perception component that is compatible with abiotic-stress signalling. Lectin receptor-like kinases (LecRLKs) are repeatedly implicated in stress responses, and genome-wide analyses coupled with expression/functional evidence support their involvement in salinity tolerance through ion homeostasis and ROS-related pathways (Xiong et al. 2024). More broadly, receptor-like kinases (RLKs), including LRR-RLKs, have emerged as key hubs connecting extracellular stress perception with downstream transcriptional and metabolic reprogramming under drought and salinity (Gandhi and Oelmüller 2023; Yan et al. 2025).

In addition to signalling components, the wsnp_Ex_c36049_44083089 region is enriched for enzymes of specialized and primary metabolism, notably multiple cytochrome P450 monooxygenases (*TraesCS2A02G005100.1, G005800.1, G005900.1, G006300.1, G006400.1*), a terpene synthase (*TraesCS2A02G006000.1*) and secondary-metabolite acyltransferases (*TraesCS2A02G006200.1, G006700.1*) (Fig. 5), gene families that together underpin the biosynthesis and modification of a wide range of specialized metabolites (terpenes, phenylpropanoids and other defense compounds) and have repeatedly been implicated in abiotic-stress acclimation, including terpene synthases and P450s that are induced by drought, salinity or oxidative stress and contribute to ROS management, cell-wall reinforcement and volatile-mediated stress signalling (Pandian et al. 2020). In this context, the co-localization of wsnp_Ex_c36049_44083089 with NLRs, LEA₂/NDR1-like receptors and candidate LecRLKs, as well as terpenoid and phenylpropanoid metabolism enzymes, provides an explanation for why this 2A locus has a strong effect on vegetation indices under rainfed conditions.

### Implications for breeding and future work

From a breeding perspective, the combination of high heritability for most spectral indices and the presence of major QTL effects as well as pleiotropic QTLs suggests that digital phenotyping can be effectively integrated into selection pipelines. The major loci on 2A, 4A, 5A, and 7A, where favorable alleles enhance greenness and pigment-retention indices under rainfed conditions, represent promising targets for MAS and genomic prediction aimed at improving drought resilience and yield stability. At the same time, several limitations need to be acknowledged in terms of QTL validation studies as well as testing their efficiency across different environments. Further, candidate-gene assignments based on physical proximity and domain annotations remain putative; integrating transcriptomic, metabolomic and functional genomic approaches will be essential to validate the roles of NLRs, LEA/NDR1-like proteins, lectin receptor-like kinases and metabolic genes in controlling spectral traits under drought.

## Conclusion

The results of this study demonstrated the synergistic effect of integrating high-throughput digital phenotyping and genome-wide SNP data using GWAS in deciphering the complex genetic architecture of vegetation indices in bread wheat. The strongest QTL effect was detected on chromosome 2A at marker wsnp_Ex_c36049_44083089 (for 17 vegetation indices), which explained a major proportion of genetic variation (up to 20%). Notably, this significant association identifies a pleiotropic locus on chromosome 2A, underscoring its potential as a target region for breeding. Further, the co-localization of candidate genes linked to stress response and metabolic processes opens new avenues for future research. Overall, our results demonstrate that GWAS of vegetation indices under rainfed conditions can reveal coherent networks of loci and candidate genes underlying canopy greenness, pigment composition, and structural traits.

## Supplementary information

The online version contains supplementary material available at the link:

## Author contributions

I.H. supervised the project and acquired funding. I.H., A.A.N., and S.R. conceptualized the research idea. A.J. and S.R. conducted the field trials. J.L., A.A.N., and S.R. performed the statistical analysis. S.R. drafted the original manuscript. I.H., J.L., A.A.N. and SR contributed to the review and editing process. All authors read and approved the final manuscript.

## Data availability

The data that support the findings of this study are available from Dr. Samira Rustamova upon request.

