## Supplementary Table S1 for "Wheat diversity reveals new genomic loci and candidate genes for vegetation indices using genome-wide association analysis"

**Table S1** List of bread wheat genotypes used in this study

| **No** | **Genotype**  **name** | **Country**  **of origin** | **Varietal type** |
| --- | --- | --- | --- |
| 1. | Gobustan | Azerbaijan | Greakum |
| 2. | Askeran | Azerbaijan | Greakum |
| 3. | Azamatli95 | Azerbaijan | Greakum |
| 4. | Tale38 | Azerbaijan | Greakum |
| 5. | Zirva85 | Azerbaijan | Erytrospermum |
| 6. | Layagatli80 | Azerbaijan | Erytrospermum |
| 7. | Gyrmyzygul1 | Azerbaijan | Erytrospermum |
| 8. | Baba75 | Azerbaijan | Erytrospermum |
| 9. | Gyzylbugda | Azerbaijan | Lutescens |
| 10. | Bezostaya1AZ | Russia | Lutescens |
| 11. | Murov2 | Azerbaijan | Lutescens |
| 12. | Azeri | Azerbaijan | Lutescens |
| 13. | Khezri | Azerbaijan | Erytrospermum |
| 14. | Yubiley90 | Azerbaijan | Erytrospermum |
| 15. | Gulustan100 | Azerbaijan | Erytrospermum |
| 16. | Fatima | Azerbaijan | Erytrospermum |
| 17. | Lider | Azerbaijan | Erytrospermum |
| 18. | Chempion | Azerbaijan | Erytrospermum |
| 19. | Matin | Azerbaijan | Lutescens |
| 20. | Ugur | Azerbaijan | Lutescens |
| 21. | Altunbashak | Azerbaijan | Erytrospermum |
| 22. | Nurlu99 | Azerbaijan | Greakum |
| 23. | Giymatli217 | Azerbaijan | Velutinum |
| 24. | Jumhuriyyat100 | Azerbaijan | Lutescens |
| 25. | Shafaq2 | Azerbaijan | Lutescens |
| 26. | Altun2 | Azerbaijan | Lutescens |
| 27. | Asad80 | Azerbaijan | Erytrospermum |
| 28. | Tunj | Azerbaijan | Erytrospermum |
| 29. | Onur | Azerbaijan | Lutescens |
| 30. | Oguz | Azerbaijan | Erytrospermum |
| 31. | Diabar | Azerbaijan | Lutescens |
| 32. | Romanna | Azerbaijan | Ferriguneum |
| 33. | Shahbughda | Azerbaijan | Hostianum |
| 34. | Kanan | Azerbaijan | Erytrospermum |
| 35. | Royal | Azerbaijan | Greakum |
| 36. | Vusal | Azerbaijan | Erytrospermum |
| 37. | Almaz | Azerbaijan | Lutescens |
| 38. | Leyla | Azerbaijan | Lutescens |
| 39. | Start | Azerbaijan | Lutescens |
| 40. | Janub | Azerbaijan | Erytrospermum |
| 41. | Vilash | Azerbaijan | Lutescens |
| 42. | Khamsa | Azerbaijan | Erytrospermum |
| 43. | Shafaq | Azerbaijan | Lutescens |
| 44. | Akinchi84 | Azerbaijan | Erytrospermum |
| 45. | Dostlug | Azerbaijan | Lutescens |
| 46. | Bayaz | Azerbaijan | Erytroleukon |
| 47. | Ruzi84 | Azerbaijan | Greakum |
| 48. | Gyrmyzybugda | Azerbaijan | Erytrospermum |
| 49. | Pirshahin | Azerbaijan | Erytrospermum |
| 50. | Gunashli | Azerbaijan | Erytrospermum |
| 51. | Parzivan1 | Azerbaijan | Erytrospermum |
| 52. | Saba | Azerbaijan | Lutescens |
| 53. | Agali | Azerbaijan | Lutescens |
| 54. | Murov | Azerbaijan | Lutescens |
| 55. | Duygu | Azerbaijan | Leukomelan |
| 56. | Barish | Azerbaijan | Greakum |
| 57. | Khirman | Azerbaijan | Greakum |
| 58. | Banu | Azerbaijan | Erytrospermum |
| 59. | Marjani | Azerbaijan | Eritroleucon |
| 60. | ID106002 | Russia | Lutescens |
| 61. | Korochenko | Russia | Lutescens |
| 62. | Bezostaya1RU | Russia | Lutescens |
| 63. | Tanya | Russia | Lutescens |
| 64. | Alekseich | Russia | Lutescens |
| 65. | Antonina | Russia | Lutescens |
| 66. | Grom | Russia | Lutescens |
| 67. | Sineva | Russia | Lutescens |
| 68. | Moskovskaya39 | Russia | Lutescens |
| 69. | Bagrat | Russia | Lutescens |
| 70. | Yumpo | Russia | Greakum |
| 71. | DF-58-03 | Russia | Lutescens |
| 72. | Astarta | Russia | Lutescens |
| 73. | Adel | Russia | Lutescens |
| 74. | Skipetr | Russia | Lutescens |
| 75. | Nemchinovskaya | Russia | Lutescens |
| 76. | Aist | Russia | Lutescens |
| 77. | Kalim | Russia | Lutescens |
| 78. | Snegurka | Russia | Lutescens |
| 79. | Lutescens982 | Russia | Lutescens |
| 80. | Moskovskaya56 | Russia | Lutescens |
| 81. | Moskovskaya40 | Russia | Lutescens |
| 82. | Saratovskaya29 | Russia | Lutescens |
| 83. | Viktoriya | Russia | Lutescens |
| 84. | Status | Russia | Lutescens |
| 85. | Niva | Russia | Lutescens |
| 86. | Ksenia | Russia | Lutescens |
| 87. | Sekletia70 | Russia | Lutescens |
| 88. | Paritet | Russia | Lutescens |
| 89. | Olimp | Russia | Lutescens |
| 90. | Bagira | Russia | Lutescens |
| 91. | Karolina | Russia | Erytrospermum |
| 92. | Stat | Russia | Lutescens |
| 93. | Firuza40 | Russia | Lutescens |
| 94. | Stavka | Russia | Lutescens |
| 95. | Arsenal | Russia | Lutescens |
| 96. | Berezit | Russia | Lutescens |
| 97. | Vassa | Russia | Lutescens |
| 98. | Neirinovka52 | Russia | Lutescens |
| 99. | Yubileynaya60 | Russia | Lutescens |
| 100. | TungesKZ | Kazakhstan | Lutescens |
| 101. | Topkapi | Türkiye | Erytrospermum |
| 102. | Garak | Türkiye | Ferriguneum |
| 103. | Mufitbey | Türkiye | Erytrospermum |
| 104. | Najibey | Türkiye | Erytrospermum |
| 105. | Korpu | Türkiye | Erytrospermum |
| 106. | Sonmaz | Türkiye | Lutescens |
| 107. | Rumeli | Türkiye | Erytrospermum |
| 108. | Karahan | Türkiye | Erytrospermum |
| 109. | Dagdash | Türkiye | Erytrospermum |
| 110. | Tigre | France | Erytrospermum |
| 111. | Midus | Austria | Erytrospermum |
| 112. | Gaudio | Austria | Erytrospermum |
| 113. | Almeria | Italy | Erytrospermum |
| 114. | Hendrix | France | Erytrospermum |
| 115. | Esperia | Italy | Erytrospermum |
| 116. | Genezis | Bulgaria | Lutescens |
| 117. | Leonida | Unknown | Erytrospermum |
| 118. | Cetic | Unknown | Erytrospermum |
| 119. | Cesiko | Unknown | Erytrospermum |
| 120. | Chayda | Unknown | Erytrospermum |
| 121. | Qualite | Italy | Erytrospermum |
| 122. | Callio | Austria | Erytrospermum |
| 123. | Monrizio | Austria | Erytrospermum |
| 124. | Dicilla | Italy | Erytrospermum |
| 125. | Renan | France | Erytrospermum |
| 126. | Barok | France | Erytrospermum |
| 127. | Balaton | Austria | Lutescens |
| 128. | Milturum1 | Gene Bank (Azerbaijan) | Milturum |
| 129. | Erythrospermum2 | Gene Bank (Azerbaijan) | Erythrospermum |
| 130. | Qraecum3 | Gene Bank (Azerbaijan) | Qraecum |
| 131. | Lutescens4 | Gene Bank (Azerbaijan) | Lutescens |
| 132. | Erythrospermum5 | Gene Bank (Azerbaijan) | Erythrospermum |
| 133. | Hostianum6 | Gene Bank (Azerbaijan) | Hostianum |
| 134. | Ferrugineum | Gene Bank (Azerbaijan) | Ferrugineum |
| 135. | Velutinum | Gene Bank (Azerbaijan) | Velutinum |
| 136. | Hostianum9 | Gene Bank (Azerbaijan) | Hostianum |
| 137. | Barbarossa10 | Gene Bank (Azerbaijan) | Barbarossa |
| 138. | Lutescens11 | Gene Bank (Azerbaijan) | Lutescens |
| 139. | Hostianum12 | Gene Bank (Azerbaijan) | Hostianum |
| 140. | Milturum13 | Gene Bank (Azerbaijan) | Milturum |
| 141. | Barbarossa14 | Gene Bank (Azerbaijan) | Barbarossa |
| 142. | Hostianum15 | Gene Bank (Azerbaijan) | Hostianum |
| 143. | Barbarossa16 | Gene Bank (Azerbaijan) | Barbarossa |
| 144. | Ferrugineum17 | Gene Bank (Azerbaijan) | Ferrugineum |
| 145. | Erythrospermum18 | Gene Bank (Azerbaijan) | Erythrospermum |
| 146. | Ferrugineum19 | Gene Bank (Azerbaijan) | Ferrugineum |
| 147. | Lutescens20 | Gene Bank (Azerbaijan) | Lutescens |
| 148. | Hostianum21 | Gene Bank (Azerbaijan) | Hostianum |
| 149. | Barbarossa22 | Gene Bank (Azerbaijan) | Barbarossa |
| 150. | Barbarossa23 | Gene Bank (Azerbaijan) | Barbarossa |
| 151. | Erythrospermum24 | Gene Bank (Azerbaijan) | Erythrospermum |
| 152. | Erythroleucon25 | Gene Bank (Azerbaijan) | Erythroleucon |
| 153. | Qraecum26 | Gene Bank (Azerbaijan) | Qraecum |
| 154. | Milturum27 | Gene Bank (Azerbaijan) | Milturum |
| 155. | Milturum28 | Gene Bank (Azerbaijan) | Milturum |
| 156. | Erythrospermum | Gene Bank (Azerbaijan) | Erythrospermum |
| 157. | Lutescens30 | Gene Bank (Azerbaijan) | Lutescens |
| 158. | Barbarossa31 | Gene Bank (Azerbaijan) | Barbarossa |
| 159. | Albidum32 | Gene Bank (Azerbaijan) | Albidum |
| 160. | Hostianum33 | Gene Bank (Azerbaijan) | Hostianum |
| 161. | Hostianum34 | Gene Bank (Azerbaijan) | Hostianum |
| 162. | Hostianum35 | Gene Bank (Azerbaijan) | Hostianum |
| 163. | Qlaucolutescens36 | Gene Bank (Azerbaijan) | Qlaucolutescens |
| 164. | Cinereum37 | Gene Bank (Azerbaijan) | Cinereum |
| 165. | Beengalense38 | Gene Bank (Azerbaijan) | Beengalense |
| 166. | Renovatum39 | Gene Bank (Azerbaijan) | Renovatum |
| 167. | Lutescens40 | Gene Bank (Azerbaijan) | Lutescens |
| 168. | Milturum41 | Gene Bank (Azerbaijan) | Milturum |
| 169. | Sardoum42 | Gene Bank (Azerbaijan) | Sardoum |
| 170. | Barbarossa43 | Gene Bank (Azerbaijan) | Barbarossa |
| 171. | Milturum44 | Gene Bank (Azerbaijan) | Milturum |
| 172. | Velutinum45 | Gene Bank (Azerbaijan) | Velutinum |
| 173. | Lutescens46 | Gene Bank (Azerbaijan) | Lutescens |
| 174. | Velutinum47 | Gene Bank (Azerbaijan) | Velutinum |
| 175. | Murinum48 | Gene Bank (Azerbaijan) | Murinum |
| 176. | Murinum49 | Gene Bank (Azerbaijan) | Murinum |
| 177. | Cianotrics50 | Gene Bank (Azerbaijan) | Cianotrics |
| 178. | Velutinum51 | Gene Bank (Azerbaijan) | Velutinum |
| 179. | Erythroleucon52 | Gene Bank (Azerbaijan) | Erythroleucon |
| 180. | Barbarossa53 | Gene Bank (Azerbaijan) | Barbarossa |
| 181. | Barbarossa54 | Gene Bank (Azerbaijan) | Barbarossa |
| 182. | Murinum55 | Gene Bank (Azerbaijan) | Murinum |
| 183. | Cianotrics56 | Gene Bank (Azerbaijan) | Cianotrics |
| 184. | Cinereum57 | Gene Bank (Azerbaijan) | Cinereum |
| 185. | Delfi58 | Gene Bank (Azerbaijan) | Delfi |
| 186. | Delfi59 | Gene Bank (Azerbaijan) | Delfi |
| 187. | Lutescenes60 | Gene Bank (Azerbaijan) | Lutescenes |
