## Supplementary Table S2 for "Wheat diversity reveals new genomic loci and candidate genes for vegetation indices using genome-wide association analysis"

**Table S2** Hydro-meteorological data of the Gobustan station for the 2022/2023 growing season

| **Months** | **Decades** | | **Air temperature, ^0^C** | | | | | **Relative air humidity (%)** | **Maximum wind speed (m/s)** | **Precipitation (mm)** | | |
| --- | --- | --- | --- | --- | --- | --- | --- | --- | --- | --- | --- | --- |
|  |  |  | **Long-term average** | | **Monthly average** | **Maximum** | **Mini-**  **mum** |  |  | **By decades** | **Monthly** | **Long-term average** |
| **2022** | | | | | | | | | | | | |
| September | | I | 19,0 | | 21,2 | 37,4 | 12,0 | 48 | 9 | 0,5 | 6,0 | 31,0 |
|  |  | II | 17,2 | | 20,2 | 30,0 | 11,0 | 49 | 18 | 2,3 |  |  |
|  |  | III | 15,2 | | 19,1 | 30,5 | 6,6 | 59 | 19 | 3,2 |  |  |
| October | | I | 13,1 | | 17,9 | 27,3 | 10,0 | 62 | 9 | -- | 21.0 | 45,0 |
|  |  | II | 11.1 | | 13,9 | 24,9 | 8,3 | 80 | 19 | 6,1 |  |  |
|  |  | III | 9,4 | | 9,9 | 18,3 | 3,8 | 95 | 13 | 14,9 |  |  |
| November | | I | 7,7 | | 7,2 | 16,2 | 1,4 | 80 | 13 | 20,1 | 81,4 | 36,0 |
|  |  | II | 5,8 | | 7,4 | 21,1 | 0,5 | 85 | 22 | 30,3 |  |  |
|  |  | III | 4.5 | | 8,5 | 18,5 | 0.5 | 83 | 13 | 31,0 |  |  |
| December | | I | 3,6 | | -0,1 | 8,4 | -3,8 | 94 | 9 | 10,0 | 11,0 | 30,0 |
|  |  | II | 1,6 | | 3,9 | 12,8 | -0,7 | 90 | 12 | 1.0 |  |  |
|  |  | III | 0,9 | | 2,3 | 7,2 | -3,4 | 87 | 18 | -- |  |  |
| **2023** | | | | | | | | | | | | |
| January | | I | 0.2 | 1,8 | | 14,4 | -7,3 | 70 | 11 | 6,2 | 28,6 | 26,0 |
|  |  | II | -0.5 | 1,1 | | 4,9 | -7,2 | 93 | 9 | 20,9 |  |  |
|  |  | III | -0.4 | 0,6 | | 6,1 | -6,9 | 87 | 9 | 1,5 |  |  |
| February | | I | -0.3 | 0,3 | | 5,9 | -5,4 | 91 | 12 | 28,0 | 43,5 | 35,0 |
|  |  | II | -0.1 | -0,3 | | 14,8 | -8,5 | 72 | 22 | 7,8 |  |  |
|  |  | III | 0.8 | 3,3 | | 17,5 | -5,3 | 48 | 25 | 7,7 |  |  |
| March | | I | 1.7 | 7,2 | | 18,4 | 0.6 | 72 | 22 | 0,7 | 4.4 | 42,0 |
|  |  | II | 2.7 | 10,7 | | 21,7 | 3,1 | 67 | 22 | 1,5 |  |  |
|  |  | III | 4.8 | 10,6 | | 22,5 | 0,3 | 63 | 19 | 2,2 |  |  |
| April | | I | 7.1 | 7,8 | | 16,9 | -0,6 | 82 | 11 | 19,7 | 35,9 | 47,0 |
|  |  | II | 9.3 | 11,0 | | 23,6 | 0,5 | 90 | 22 | 13,0 |  |  |
|  |  | III | 11.2 | 16,1 | | 27,1 | 7,4 | 67 | 11 | 3,2 |  |  |
| May | | I | 13.1 | 13,0 | | 23,1 | 6,4 | 77 | 10 | 60,7 | 68,3 | 47,0 |
|  |  | II | 15.1 | 12,7 | | 23,8 | 5,7 | 74 | 9 | 4,9 |  |  |
|  |  | III | 16.6 | 21,8 | | 33,6 | 12,9 | 57 | 10 | 2,7 |  |  |
| June | | I | 18.1 | 20,4 | | 32,0 | 11,7 | 64 | 10 | 2,4 | 4,1 | 40,0 |
|  |  | II | 19.6 | 23,4 | | 32,8 | 15,0 | 68 | 10 | 1,2 |  |  |
|  |  | III | 20.7 | 21,1 | | 32,1 | 10,8 | 62 | 10 | 0,5 |  |  |
| July | | I | 21.9 | 24,2 | | 36,8 | 13,0 | 66 | 10 | 10,3 | 31,0 | 14,0 |
|  |  | II | 23.1 | 21,5 | | 31,6 | 13,4 | 73 | 12 | 20,7 |  |  |
|  |  | III | 22.9 | 25,4 | | 35,9 | 16,1 | 66 | 11 | -- |  |  |
| August | | I | 22.7 | 24,8 | | 36,1 | 19,0 | 61 | 11 | -- | 21,1 | 13,0 |
|  |  | II | 22.4 | 25,7 | | 37,1 | 16,4 | 64 | 11 | 0,8 |  |  |
|  |  | III | 20.8 | 22.2 | | 34,9 | 15,3 | 77 | 11 | 20,3 |  |  |
| **Average:** | | | 10,63 | 12,7 | |  | | | | **Total:** | 356,3 | 406,0 |
