## Supplementary Table S3 for "Wheat diversity reveals new genomic loci and candidate genes for vegetation indices using genome-wide association analysis"

### Table S3 Multispectral vegetation indices used in the study

| **Abbreviation** | **Name** | **Equation** | **Reference** |
| --- | --- | --- | --- |
| NDVI | Normalized Difference Vegetation Index | NDVI = (R800 – R670) / (R800 + R670) | Rouse et al. (1974) |
| SR | Simple Ratio Index | SR = R800 / R670 | Jordan (1969)  Rouse et al. (1974) |
| MCARI1 | Modified Chlorophyll Absorption in Reflectance Index 1 | MCARI1 = 1.2 × [2.5 × (R750 − R670) − 1.3 × (R700 − R550)] | Haboudane et al. (2002) |
| OSAVI | Optimized Soil-Adjusted Vegetation Index | OSAVI = (1 + 0.16) × (R790 − R670) / (R790 + R670 + 0.16) | Rondeaux et al. (1996) |
| G | Greenness Index | G = R554 / R677 | Gitelson et al. (1996) |
| MCARI | Modified Chlorophyll Absorption in Reflectance Index | MCARI = [(R700 − R670) − 0.2 × (R700 − R550)] × (R700 / R670) | Daughtry et al. (2000) |
| TCARI | Transformed Chlorophyll Absorption in Reflectance Index | TCARI = 3 × [(R700 − R670) − 0.2 × (R700 − R550) × (R700 / R670)] | Haboudane et al. (2002) |
| TVI | Triangular Vegetation Index | TVI = 0.5 × [120 × (R750 − R550) − 200 × (R670 − R550)] | Broge & Leblanc (2001) |
| ZMI | Zarco-Tejada & Miller Index | ZMI = R750 / R710 | Zarco-Tejada et al. (2001) |
| SPRI | Simple Pigment Ratio Index | SPRI = R430 / R680 | Peñuelas et al. (1995) |
| NPQI | Normalized Phaeophytinization Index | NPQI = (R415 − R435) / (R415 + R435) | Barnes et al. (1992) |
| PRI | Photochemical Reflectance Index | PRI = (R531 − R570) / (R531 + R570) | Gamon et al. (1997) |
| NPCI | Normalized Pigment Chlorophyll Index | NPCl = (R680 − R430) / (R680 + R430) | Peñuelas et al. (1995) |
| Ctr1 | Carter Index 1 | Ctrl = R695 / R420 | Carter (1994) |
| Ctr2 | Carter Index 2 | Ctr2 = R695 / R760 | Carter (1994) |
| Lic1 | Lichtenthaler Index 1 | Lic1 = (R790 − R680) / (R790 + R680) | Lichtenthaler et al. (1996) |
| Lic2 | Lichtenthaler Index 2 | Lic2 = R440 / R690 | Lichtenthaler et al. (1996) |
| SIPI | Structure Insensitive Pigment Index | SIPI = (R800 − R445) / (R800 + R680) | Peñuelas & Filella (1998) |
| GM1 | Gitelson & Merzlyak Index 1 | GM1 = R750 / R550 | Gitelson & Merzlyak (1994) |
| GM2 | Gitelson & Merzlyak Index 2 | GM2 = R750 / R700 | Gitelson & Merzlyak (1997) |
| ARI1 | Anthocyanin Reflectance Index 1 | ARI1 = (1 / R550) − (1 / R700) | Gitelson et al. (2001) |
| ARI2 | Anthocyanin Reflectance Index 2 | ARI2 = (R800 / R550) − (R800 / R700) | Gitelson et al. (2001) |
| CRI1 | Carotenoid Reflectance Index 1 | CRI1 = (1 / R510) − (1 / R550) | Gitelson et al. (2002) |
| CRI2 | Carotenoid Reflectance Index 2 | CRI2 = (1 / R510) − (1 / R700) | Gitelson et al. (2002) |
| RDVI | Renormalized Difference Vegetation Index | RDVI = (R800 − R670) / √(R800 + R670) | Roujean & Breon (1995) |
