## Supplementary Table S5 for "Wheat diversity reveals new genomic loci and candidate genes for vegetation indices using genome-wide association analysis"

**Table S4** LD-independent lead SNPs surpassing the Bonferroni threshold for spectral vegetation indices

| **Trait** | **Marker** | **Chr** | **pos** | **MAF** | **major_allel** | **Minor_Allel** | **F_value** | **ProbF** | **FDR_P** | **Explained_Var** | **Mean_A1A1** | **Mean_A2A2** |
| --- | --- | --- | --- | --- | --- | --- | --- | --- | --- | --- | --- | --- |
| NDVI | wsnp_Ex_c36049_44083089 | 2A | 2925147 | 0,19672131 | C | T | 46,9477061 | 1,0489E-10 | 1,9187E-06 | 20,0834304 | 0,528777 | 0,26545018 |
| NDVI | IAAV3697 | 4A | 38335364 | 0,17647059 | T | C | 44,449408 | 2,9163E-10 | 1,9187E-06 | 19,3722042 | 0,52702859 | 0,2601421 |
| NDVI | AX-158584871 | 5A | 535796991 | 0,2311828 | A | G | 42,6079297 | 6,246E-10 | 3,082E-06 | 18,3148524 | 0,53680679 | 0,30159906 |
| NDVI | wsnp_Ex_c7216_12390889 | 7A | 524972333 | 0,34054054 | A | G | 41,6879612 | 9,1615E-10 | 3,6164E-06 | 18,3839298 | 0,551101 | 0,34082762 |
| NDVI | BS00086534_51 | 2D | 656028039 | 0,33125 | G | A | 40,235171 | 1,6835E-09 | 5,538E-06 | 14,2613749 | 0,5557054 | 0,35625486 |
| NDVI | AX-158584345 | 5A | 524940962 | 0,25225225 | A | G | 36,8585583 | 7,0454E-09 | 1,9865E-05 | 6,17563201 | 0,52614893 | 0,60525824 |
| NDVI | Excalibur_rep_c68688_103 | 5D | 45176107 | 0,14438503 | T | C | 36,3846258 | 8,6302E-09 | 2,1292E-05 | 16,4350283 | 0,51842736 | 0,25180426 |
| NDVI | Ra_c108731_554 | 5B | 61846046 | 0,17741935 | G | T | 33,0819839 | 3,5991E-08 | 6,4577E-05 | 15,112561 | 0,52073314 | 0,28488774 |
| NDVI | RAC875_rep_c111494_195 | 1B | 665826070 | 0,42162162 | T | C | 32,5836967 | 4,4741E-08 | 7,3002E-05 | 14,9366224 | 0,40203169 | 0,58390398 |
| NDVI | AX-95208179 | 1D | 335597569 | 0,26344086 | C | T | 32,0828166 | 5,5716E-08 | 7,8547E-05 | 14,7235776 | 0,53207492 | 0,33018779 |
| NDVI | IAAV149 | 1D | 429717285 | 0,14673913 | A | G | 31,782643 | 6,3561E-08 | 8,1555E-05 | 14,5110945 | 0,51639154 | 0,2655165 |
| NDVI | AX-158545574 | 1D | 455784529 | 0,38502674 | T | C | 30,708927 | 1,0201E-07 | 0,00010203 | 14,2362801 | 0,54894357 | 0,36970252 |
| NDVI | RAC875_c64377_732 | 2D | 573933688 | 0,40437158 | C | T | 30,6500421 | 1,047E-07 | 0,00010203 | 14,0567114 | 0,40455111 | 0,58294258 |
| NDVI | AX-158569026 | 1A | 586999805 | 0,32795699 | T | C | 30,4629423 | 1,1373E-07 | 0,00010203 | 14,0094918 | 0,53890136 | 0,35414247 |
| NDVI | AX-95168570 | 2B | 812643564 | 0,43646409 | G | C | 30,3276061 | 1,2075E-07 | 0,00010362 | 13,8029771 | 0,55438583 | 0,37843578 |
| NDVI | RAC875_c33083_451 | 3B | 841569821 | 0,26344086 | A | G | 29,9636587 | 1,4189E-07 | 0,00011555 | 13,8866058 | 0,53054092 | 0,33447675 |
| NDVI | IAAV912 | 7A | 20370858 | 0,28877005 | G | T | 29,8937657 | 1,4636E-07 | 0,00011555 | 13,9109506 | 0,53486763 | 0,34462401 |
| NDVI | Kukri_rep_c102102_273 | 1A | 571167863 | 0,40322581 | C | T | 29,6355449 | 1,6415E-07 | 0,00012461 | 13,7035229 | 0,4079393 | 0,58282326 |
| NDVI | AX-95099091 | 5B | 424325426 | 0,13903743 | A | G | 29,0371194 | 2,1427E-07 | 0,0001411 | 13,5663942 | 0,51414615 | 0,26806009 |
| NDVI | AX-158542802 | 5A | 458637478 | 0,14606742 | A | C | 29,0350805 | 2,1446E-07 | 0,0001411 | 13,299583 | 0,51255911 | 0,26806009 |
| NDVI | BobWhite_c16397_524 | 5D | 498621954 | 0,49462366 | T | G | 27,3698164 | 4,5236E-07 | 0,00022713 | 12,8752273 | 0,56167645 | 0,39533486 |
| NDVI | BobWhite_c30138_69 | 7D | 20933108 | 0,34224599 | T | C | 27,3311155 | 4,6031E-07 | 0,00022713 | 12,8719314 | 0,53975451 | 0,36495761 |
| NDVI | AX-111457218 | 4B | 533851398 | 0,33879781 | T | C | 27,2213824 | 4,8365E-07 | 0,00023282 | 12,2191419 | 0,54260172 | 0,37025875 |
| NDVI | Kukri_c13045_302 | 5D | 244100832 | 0,44385027 | G | T | 26,3952843 | 7,0248E-07 | 0,00027738 | 12,4862219 | 0,55290204 | 0,38849735 |
| NDVI | TA003390-0807 | 2A | 787686566 | 0,46236559 | A | C | 25,3334222 | 1,1381E-06 | 0,00039025 | 12,0262415 | 0,5538087 | 0,39259901 |
| NDVI | IAAV4569 | 1B | 582731648 | 0,1657754 | G | A | 24,9693276 | 1,3438E-06 | 0,00042075 | 11,8918929 | 0,51546602 | 0,30110947 |
| NDVI | JD_c1555_151 | 6B | 574415823 | 0,2513369 | G | T | 23,8084337 | 2,2881E-06 | 0,00064515 | 11,402046 | 0,52515707 | 0,34521494 |
| NDVI | RAC875_c18659_402 | 6D | 32298303 | 0,12365591 | A | G | 23,6307026 | 2,4832E-06 | 0,0006903 | 11,3242491 | 0,5089865 | 0,27204837 |
| ARI1 | GENE-0511_161 | 1A | 512571385 | 0,47252747 | A | G | 26,8629631 | 5,6854E-07 | 0,00335656 | 12,6659451 | -0,06067724 | -0,19910803 |
| ARI1 | BS00021702_51 | 1D | 416991781 | 0,49197861 | T | C | 26,4974784 | 6,7071E-07 | 0,00335656 | 12,528508 | -0,05864072 | -0,19428486 |
| ARI2 |  |  |  |  |  |  |  |  |  |  |  |  |
| CRI1 | IAAV3697 | 4A | 38335364 | 0,17647059 | T | C | 34,7228509 | 1,7649E-08 | 0,00017417 | 15,8030222 | 1,987148 | 0,90200282 |
| CRI1 | wsnp_Ex_c7216_12390889 | 7A | 524972333 | 0,34054054 | A | G | 32,778006 | 4,1098E-08 | 0,00022108 | 14,6133021 | 2,07155135 | 1,2279291 |
| CRI1 | wsnp_Ex_c36049_44083089 | 2A | 2925147 | 0,19672131 | C | T | 32,5804403 | 4,4805E-08 | 0,00022108 | 14,8840811 | 1,9859676 | 0,96538818 |
| CRI1 | BS00086534_51 | 2D | 656028039 | 0,33125 | G | A | 31,4897327 | 7,2296E-08 | 0,00028538 | 12,0555851 | 2,13278205 | 1,31486076 |
| CRI1 | AX-95168570 | 2B | 812643564 | 0,43646409 | G | C | 28,1468885 | 3,1904E-07 | 0,00089956 | 13,0275438 | 2,13143811 | 1,36165006 |
| CRI1 | AX-158569026 | 1A | 586999805 | 0,32795699 | T | C | 25,9077144 | 8,7635E-07 | 0,00216208 | 12,2547987 | 2,04707888 | 1,26907572 |
| CRI1 | IAAV149 | 1D | 429717285 | 0,14673913 | A | G | 25,0331028 | 1,3052E-06 | 0,00257609 | 11,7113473 | 1,94073947 | 0,92619571 |
| CRI1 | TA003390-0807 | 2A | 787686566 | 0,46236559 | A | C | 24,7260601 | 1,5019E-06 | 0,00269476 | 11,7608193 | 2,12368477 | 1,40602927 |
| CRI1 | AX-158584871 | 5A | 535796991 | 0,2311828 | A | G | 24,2907324 | 1,8334E-06 | 0,00280861 | 11,3550794 | 1,99961245 | 1,16604446 |
| CRI1 | Excalibur_rep_c68688_103 | 5D | 45176107 | 0,14438503 | T | C | 24,2711198 | 1,8499E-06 | 0,00280861 | 11,5979308 | 1,9412331 | 0,93294769 |
| CRI2 | IAAV3697 | 4A | 38335364 | 0,17647059 | T | C | 32,875329 | 3,9388E-08 | 0,0003887 | 15,0890553 | 1,85429222 | 0,81236298 |
| CRI2 | wsnp_Ex_c36049_44083089 | 2A | 2925147 | 0,19672131 | C | T | 26,630058 | 6,3166E-07 | 0,00261438 | 12,4624849 | 1,84067992 | 0,92308671 |
| CRI2 | BS00086534_51 | 2D | 656028039 | 0,33125 | G | A | 26,566386 | 6,5012E-07 | 0,00261438 | 10,2378088 | 1,98225196 | 1,24453196 |
| CRI2 | wsnp_Ex_c7216_12390889 | 7A | 524972333 | 0,34054054 | A | G | 26,1229862 | 7,9476E-07 | 0,00261438 | 12,3598095 | 1,93345474 | 1,17080173 |
| CRI2 | AX-95168570 | 2B | 812643564 | 0,43646409 | G | C | 24,2266994 | 1,888E-06 | 0,00532341 | 11,4193472 | 1,98414074 | 1,27601232 |
| ctr1 | wsnp_Ex_c5690_9994305 | 4A | 38692608 | 0,48924731 | G | A | 24,4662622 | 1,69E-06 | 0,01772222 | 11,6656112 | 1,28393016 | 1,12974621 |
| Ctr2 | Excalibur_rep_c68688_103 | 5D | 45176107 | 0,14438503 | T | C | 44,0509135 | 3,4369E-10 | 6,7834E-06 | 19,2319309 | 0,37707745 | 0,73720598 |
| Ctr2 | wsnp_Ex_c36049_44083089 | 2A | 2925147 | 0,19672131 | C | T | 41,3648859 | 1,0485E-09 | 1,0347E-05 | 18,135893 | 0,37113226 | 0,68358019 |
| Ctr2 | AX-158584871 | 5A | 535796991 | 0,2311828 | A | G | 39,0853871 | 2,7335E-09 | 1,7984E-05 | 17,1537924 | 0,36063402 | 0,64487645 |
| Ctr2 | wsnp_Ex_c7216_12390889 | 7A | 524972333 | 0,34054054 | A | G | 37,0654311 | 6,4492E-09 | 2,9164E-05 | 16,6807743 | 0,34457336 | 0,59466782 |
| Ctr2 | IAAV3697 | 4A | 38335364 | 0,17647059 | T | C | 36,3218218 | 8,8658E-09 | 2,9164E-05 | 16,4113152 | 0,37494753 | 0,68166769 |
| Ctr2 | Ra_c108731_554 | 5B | 61846046 | 0,17741935 | G | T | 33,6049268 | 2,866E-08 | 8,0808E-05 | 15,3179683 | 0,3777367 | 0,6742161 |
| Ctr2 | IAAV149 | 1D | 429717285 | 0,14673913 | A | G | 32,3928962 | 4,8637E-08 | 0,00011999 | 14,7725066 | 0,38325377 | 0,69931125 |
| Ctr2 | BS00086534_51 | 2D | 656028039 | 0,33125 | G | A | 31,9332817 | 5,9493E-08 | 0,00013047 | 12,0388605 | 0,34246534 | 0,57135301 |
| Ctr2 | AX-95208179 | 1D | 335597569 | 0,26344086 | C | T | 29,6513487 | 1,63E-07 | 0,00026809 | 13,764771 | 0,36612785 | 0,60986389 |
| Ctr2 | RAC875_c33083_451 | 3B | 841569821 | 0,26344086 | A | G | 27,8917698 | 3,5773E-07 | 0,00040813 | 13,0549544 | 0,36780558 | 0,6051731 |
| Ctr2 | RAC875_rep_c111494_195 | 1B | 665826070 | 0,42162162 | T | C | 27,6372381 | 4,0107E-07 | 0,00040813 | 12,969071 | 0,51970023 | 0,3080938 |
| Ctr2 | AX-94824007 | 2B | 358483 | 0,09677419 | A | G | 27,5690089 | 4,1357E-07 | 0,00040813 | 12,9546286 | 0,39426221 | 0,74658029 |
| Ctr2 | AX-158545574 | 1D | 455784529 | 0,38502674 | T | C | 27,2047075 | 4,873E-07 | 0,00043717 | 12,8200302 | 0,34730194 | 0,55968375 |
| Ctr2 | AX-158569026 | 1A | 586999805 | 0,32795699 | T | C | 26,3187733 | 7,2725E-07 | 0,00055821 | 12,3514777 | 0,35996513 | 0,57657834 |
| Ctr2 | JD_c1555_151 | 6B | 574415823 | 0,2513369 | G | T | 26,0620193 | 8,1706E-07 | 0,00055821 | 12,3480384 | 0,37030806 | 0,60412394 |
| Ctr2 | AX-158582092 | 4B | 611116205 | 0,22994652 | T | C | 25,9695939 | 8,5207E-07 | 0,00055821 | 12,3096383 | 0,3737368 | 0,61439194 |
| Ctr2 | RAC875_c64377_732 | 2D | 573933688 | 0,40437158 | C | T | 25,6861888 | 9,692E-07 | 0,00055821 | 12,0650538 | 0,51633995 | 0,30998284 |
| Ctr2 | IAAV4569 | 1B | 582731648 | 0,1657754 | G | A | 25,6397765 | 9,8988E-07 | 0,00055821 | 12,1723337 | 0,38418438 | 0,65497384 |
| Ctr2 | AX-158584345 | 5A | 524940962 | 0,25225225 | A | G | 25,2870796 | 1,1624E-06 | 0,00060597 | 4,75818755 | 0,38097455 | 0,28901864 |
| Ctr2 | RAC875_c18659_402 | 6D | 32298303 | 0,12365591 | A | G | 25,2672121 | 1,173E-06 | 0,00060597 | 12,0098323 | 0,39190831 | 0,69657769 |
| Ctr2 | AX-95168570 | 2B | 812643564 | 0,43646409 | G | C | 25,2220233 | 1,1974E-06 | 0,00060597 | 11,8100706 | 0,34327864 | 0,54648614 |
| Ctr2 | Kukri_rep_c102102_273 | 1A | 571167863 | 0,40322581 | C | T | 24,8193626 | 1,4391E-06 | 0,00065869 | 11,7384496 | 0,51242132 | 0,31032389 |
| Ctr2 | BobWhite_c11503_296 | 6D | 458697272 | 0,2459893 | G | A | 24,7790481 | 1,4659E-06 | 0,00065869 | 11,8119747 | 0,37241456 | 0,60274999 |
| Ctr2 | AX-95099091 | 5B | 424325426 | 0,13903743 | A | G | 24,739279 | 1,4928E-06 | 0,00065869 | 11,7952532 | 0,3892388 | 0,67575025 |
| Ctr2 | AX-158542802 | 5A | 458637478 | 0,14606742 | A | C | 24,6418977 | 1,5608E-06 | 0,00065869 | 11,5518733 | 0,39124067 | 0,67575025 |
| Ctr2 | BobWhite_c16397_524 | 5D | 498621954 | 0,49462366 | T | G | 24,6311435 | 1,5685E-06 | 0,00065869 | 11,7369717 | 0,33168833 | 0,52999303 |
| G | BS00086534_51 | 2D | 656028039 | 0,33125 | G | A | 42,5115848 | 6,5012E-10 | 1,0111E-05 | 15,0798513 | 1,47617881 | 1,23062874 |
| G | IAAV3697 | 4A | 38335364 | 0,17647059 | T | C | 40,4521532 | 1,5369E-09 | 1,0111E-05 | 17,9426777 | 1,43577136 | 1,12783362 |
| G | wsnp_Ex_c7216_12390889 | 7A | 524972333 | 0,34054054 | A | G | 37,5195884 | 5,3132E-09 | 2,6217E-05 | 16,8221544 | 1,46441278 | 1,22326687 |
| G | wsnp_Ex_c36049_44083089 | 2A | 2925147 | 0,19672131 | C | T | 35,5652563 | 1,2271E-08 | 4,8439E-05 | 15,8829071 | 1,43204177 | 1,15141527 |
| G | Excalibur_c24638_380 | 5D | 359180839 | 0,12834225 | A | G | 32,4869811 | 4,6675E-08 | 0,0001316 | 14,9374371 | 1,4225297 | 1,10228994 |
| G | AX-109816649 | 5A | 521757324 | 0,34234234 | T | C | 32,0044375 | 5,7664E-08 | 0,00014227 | 5,57455433 | 1,4859938 | 1,38951984 |
| G | AX-95168570 | 2B | 812643564 | 0,43646409 | G | C | 31,0447943 | 8,7951E-08 | 0,00019288 | 14,1134733 | 1,47398951 | 1,26061031 |
| G | AX-158584871 | 5A | 535796991 | 0,2311828 | A | G | 30,6098795 | 1,0657E-07 | 0,00021034 | 13,8809122 | 1,44121667 | 1,19575042 |
| G | AX-94791713 | 7A | 19958691 | 0,36898396 | T | C | 28,1257636 | 3,2208E-07 | 0,00033457 | 13,1967919 | 1,45841558 | 1,24977189 |
| G | TA003390-0807 | 2A | 787686566 | 0,46236559 | A | C | 27,2523258 | 4,7695E-07 | 0,00042537 | 12,8117888 | 1,47271589 | 1,27323182 |
| G | TG0028 | 5D | 3622961 | 0,48648649 | G | A | 27,0724226 | 5,1725E-07 | 0,00042537 | 12,7026093 | 1,47983266 | 1,28117889 |
| G | RAC875_c18659_402 | 6D | 32298303 | 0,12365591 | A | G | 26,662624 | 6,2242E-07 | 0,00049139 | 12,5963508 | 1,41836073 | 1,11876429 |
| G | Ra_c108731_554 | 5B | 61846046 | 0,17741935 | G | T | 26,3717866 | 7,0999E-07 | 0,0005082 | 12,4454636 | 1,42593859 | 1,16934144 |
| G | IAAV149 | 1D | 429717285 | 0,14673913 | A | G | 26,3379576 | 7,2096E-07 | 0,0005082 | 12,2954319 | 1,42129175 | 1,14443715 |
| G | AX-158542802 | 5A | 458637478 | 0,14606742 | A | C | 25,7491886 | 9,4182E-07 | 0,00064099 | 11,9194937 | 1,41928829 | 1,1417114 |
| G | AX-95099091 | 5B | 424325426 | 0,13903743 | A | G | 25,4246744 | 1,0917E-06 | 0,00071825 | 12,0825538 | 1,42014163 | 1,1417114 |
| G | AX-111584542 | UN | 222960893 | 0,12903226 | T | C | 25,21818 | 1,1995E-06 | 0,00076368 | 11,8979308 | 1,41647145 | 1,13060414 |
| G | AX-158545574 | 1D | 455784529 | 0,38502674 | T | C | 24,8118763 | 1,4441E-06 | 0,00081432 | 11,8257731 | 1,45683909 | 1,26098338 |
| G | BobWhite_c30138_69 | 7D | 20933108 | 0,34224599 | T | C | 24,6601661 | 1,5478E-06 | 0,00082567 | 11,7619701 | 1,44998967 | 1,24966515 |
| G | BS00018740_51 | 5D | 542233108 | 0,42162162 | T | C | 24,0297122 | 2,0668E-06 | 0,00097124 | 11,1130569 | 1,4631542 | 1,27512362 |
| GM1 | wsnp_Ex_c36049_44083089 | 2A | 2925147 | 0,19672131 | C | T | 40,8183258 | 1,318E-09 | 2,6014E-05 | 18,016302 | 2,82840382 | 1,94759372 |
| GM1 | wsnp_Ex_c7216_12390889 | 7A | 524972333 | 0,34054054 | A | G | 35,3819558 | 1,3279E-08 | 0,00013104 | 15,970726 | 2,89370185 | 2,20173752 |
| GM1 | AX-158584345 | 5A | 524940962 | 0,25225225 | A | G | 34,0922901 | 2,3191E-08 | 0,00015258 | 6,83083956 | 2,77948534 | 3,1747603 |
| GM1 | IAAV3697 | 4A | 38335364 | 0,17647059 | T | C | 32,6508748 | 4,3446E-08 | 0,0001715 | 15,0014903 | 2,80829575 | 1,97904276 |
| GM1 | AX-158584871 | 5A | 535796991 | 0,2311828 | A | G | 31,8747236 | 6,1042E-08 | 0,0002008 | 14,429351 | 2,84047199 | 2,10333799 |
| GM1 | AX-95208179 | 1D | 335597569 | 0,26344086 | C | T | 29,8866049 | 1,4682E-07 | 0,00034408 | 13,8838942 | 2,84181835 | 2,14958018 |
| GM1 | IAAV149 | 1D | 429717285 | 0,14673913 | A | G | 29,6969573 | 1,5973E-07 | 0,00034408 | 13,7434019 | 2,78788932 | 1,92582445 |
| GM1 | RAC875_rep_c111494_195 | 1B | 665826070 | 0,42162162 | T | C | 29,5654301 | 1,6935E-07 | 0,00034408 | 13,6386989 | 2,39537095 | 3,00885151 |
| GM1 | AX-158569026 | 1A | 586999805 | 0,32795699 | T | C | 29,4318921 | 1,7971E-07 | 0,00034408 | 13,5276062 | 2,86496245 | 2,22399066 |
| GM1 | RAC875_c64377_732 | 2D | 573933688 | 0,40437158 | C | T | 28,7907119 | 2,3918E-07 | 0,00034408 | 13,1843383 | 2,39893889 | 3,00844517 |
| GM1 | BS00086534_51 | 2D | 656028039 | 0,33125 | G | A | 28,7013042 | 2,4893E-07 | 0,00034408 | 11,0026567 | 2,89662941 | 2,27798556 |
| GM1 | RAC875_c33083_451 | 3B | 841569821 | 0,26344086 | A | G | 28,3818058 | 2,8717E-07 | 0,00035424 | 13,2778175 | 2,83779328 | 2,16083395 |
| GM1 | Kukri_rep_c102102_273 | 1A | 571167863 | 0,40322581 | C | T | 27,1363093 | 5,0256E-07 | 0,00052205 | 12,6833051 | 2,41689357 | 3,01093779 |
| GM1 | BobWhite_c30138_69 | 7D | 20933108 | 0,34224599 | T | C | 26,7887111 | 5,8794E-07 | 0,00055258 | 12,6487908 | 2,87134774 | 2,25953413 |
| GM1 | Ra_c108731_554 | 5B | 61846046 | 0,17741935 | G | T | 26,270296 | 7,4341E-07 | 0,00063351 | 12,4128236 | 2,79335746 | 2,03863184 |
| GM1 | Kukri_c16352_687 | 3D | 611641089 | 0,17112299 | G | A | 25,951296 | 8,5918E-07 | 0,00063351 | 12,302032 | 2,79203178 | 2,03190722 |
| GM1 | IAAV912 | 7A | 20370858 | 0,28877005 | G | T | 25,6603697 | 9,8065E-07 | 0,00064517 | 12,1809193 | 2,84346954 | 2,21489829 |
| GM1 | Kukri_c13045_302 | 5D | 244100832 | 0,44385027 | G | T | 25,0899111 | 1,2718E-06 | 0,00071718 | 11,9424636 | 2,91393631 | 2,34622386 |
| GM1 | AX-158545574 | 1D | 455784529 | 0,38502674 | T | C | 24,9917355 | 1,3301E-06 | 0,00072923 | 11,9012948 | 2,88475433 | 2,30610012 |
| GM1 | Excalibur_c24638_380 | 5D | 359180839 | 0,12834225 | A | G | 24,764806 | 1,4755E-06 | 0,00075289 | 11,8059871 | 2,76956845 | 1,93109579 |
| GM1 | IAAV4569 | 1B | 582731648 | 0,1657754 | G | A | 24,5117662 | 1,6567E-06 | 0,00075289 | 11,69947 | 2,78640765 | 2,03568914 |
| GM1 | BobWhite_c16397_524 | 5D | 498621954 | 0,49462366 | T | G | 24,4154934 | 1,7314E-06 | 0,00075289 | 11,6422352 | 2,93593449 | 2,37744002 |
| GM1 | AX-111457218 | 4B | 533851398 | 0,33879781 | T | C | 24,1377699 | 1,9667E-06 | 0,00080868 | 11,0057796 | 2,86639683 | 2,28836857 |
| GM1 | AX-158542802 | 5A | 458637478 | 0,14606742 | A | C | 23,6786172 | 2,429E-06 | 0,00091985 | 10,9464306 | 2,7647634 | 1,98174034 |
| GM2 | AX-158584345 | 5A | 524940962 | 0,25225225 | A | G | 30,2070894 | 1,2737E-07 | 0,00251392 | 8,45200228 | 2,84654432 | 3,5343934 |
| GM2 | wsnp_Ex_c7216_12390889 | 7A | 524972333 | 0,34054054 | A | G | 27,3908901 | 4,4809E-07 | 0,00360223 | 12,886669 | 3,03629195 | 2,24687235 |
| GM2 | wsnp_Ex_c36049_44083089 | 2A | 2925147 | 0,19672131 | C | T | 26,9463421 | 5,4753E-07 | 0,00360223 | 12,6992301 | 2,94756572 | 2,00833889 |
| GM2 | TA011876-1039 | 3D | 612524357 | 0,17204301 | A | G | 23,6454104 | 2,4665E-06 | 0,01014122 | 7,8263357 | 2,85632309 | 2,09417908 |
| Lic1 | wsnp_Ex_c36049_44083089 | 2A | 2925147 | 0,19672131 | C | T | 47,1225599 | 9,7691E-11 | 1,7135E-06 | 20,1049784 | 0,55052321 | 0,26039452 |
| Lic1 | IAAV3697 | 4A | 38335364 | 0,17647059 | T | C | 44,7242106 | 2,6045E-10 | 1,7135E-06 | 19,4686535 | 0,54908268 | 0,25442997 |
| Lic1 | AX-158584871 | 5A | 535796991 | 0,2311828 | A | G | 42,8435737 | 5,6639E-10 | 2,3006E-06 | 18,3772646 | 0,55988983 | 0,30041844 |
| Lic1 | wsnp_Ex_c7216_12390889 | 7A | 524972333 | 0,34054054 | A | G | 42,7747369 | 5,828E-10 | 2,3006E-06 | 18,7742956 | 0,57637748 | 0,3423571 |
| Lic1 | BS00086534_51 | 2D | 656028039 | 0,33125 | G | A | 42,2258499 | 7,3216E-10 | 2,4084E-06 | 14,7565706 | 0,58235897 | 0,35901686 |
| Lic1 | Excalibur_rep_c68688_103 | 5D | 45176107 | 0,14438503 | T | C | 37,1737649 | 6,1577E-09 | 1,7362E-05 | 16,7318427 | 0,53986231 | 0,24359084 |
| Lic1 | AX-158584345 | 5A | 524940962 | 0,25225225 | A | G | 36,4713545 | 8,3154E-09 | 2,0515E-05 | 6,01569348 | 0,54932034 | 0,63106194 |
| Lic1 | Ra_c108731_554 | 5B | 61846046 | 0,17741935 | G | T | 33,4475477 | 3,0691E-08 | 5,5068E-05 | 15,2540624 | 0,54223655 | 0,28128682 |
| Lic1 | RAC875_rep_c111494_195 | 1B | 665826070 | 0,42162162 | T | C | 32,9105366 | 3,8787E-08 | 5,8887E-05 | 15,0729019 | 0,41107363 | 0,61228555 |
| Lic1 | AX-95208179 | 1D | 335597569 | 0,26344086 | C | T | 31,7435845 | 6,4661E-08 | 7,3761E-05 | 14,5907671 | 0,55424718 | 0,33291416 |
| Lic1 | AX-95168570 | 2B | 812643564 | 0,43646409 | G | C | 31,6535939 | 6,727E-08 | 7,3761E-05 | 14,3217167 | 0,58094405 | 0,38354613 |
| Lic1 | AX-158545574 | 1D | 455784529 | 0,38502674 | T | C | 31,1344971 | 8,454E-08 | 8,1937E-05 | 14,4051493 | 0,57353812 | 0,37497274 |
| Lic1 | IAAV149 | 1D | 429717285 | 0,14673913 | A | G | 31,0647283 | 8,7181E-08 | 8,1937E-05 | 14,2189577 | 0,536736 | 0,26324394 |
| Lic1 | RAC875_c64377_732 | 2D | 573933688 | 0,40437158 | C | T | 30,8119214 | 9,747E-08 | 8,5518E-05 | 14,1397724 | 0,4141497 | 0,61121582 |
| Lic1 | IAAV912 | 7A | 20370858 | 0,28877005 | G | T | 30,4189338 | 1,1597E-07 | 9,1552E-05 | 14,1208264 | 0,55804154 | 0,34695178 |
| Lic1 | AX-158569026 | 1A | 586999805 | 0,32795699 | T | C | 29,8453489 | 1,4954E-07 | 0,00010577 | 13,7762384 | 0,5615539 | 0,35977743 |
| Lic1 | AX-95099091 | 5B | 424325426 | 0,13903743 | A | G | 29,8390825 | 1,4996E-07 | 0,00010577 | 13,8890383 | 0,53521174 | 0,26099353 |
| Lic1 | Kukri_rep_c102102_273 | 1A | 571167863 | 0,40322581 | C | T | 29,8376725 | 1,5005E-07 | 0,00010577 | 13,790349 | 0,41760439 | 0,61081509 |
| Lic1 | AX-158542802 | 5A | 458637478 | 0,14606742 | A | C | 29,7328507 | 1,572E-07 | 0,00010699 | 13,5904761 | 0,53316363 | 0,26099353 |
| Lic1 | RAC875_c33083_451 | 3B | 841569821 | 0,26344086 | A | G | 29,5743318 | 1,6868E-07 | 0,00011097 | 13,731108 | 0,55250316 | 0,3377903 |
| Lic1 | BobWhite_c16397_524 | 5D | 498621954 | 0,49462366 | T | G | 27,6019326 | 4,0749E-07 | 0,00020298 | 12,9729154 | 0,58752019 | 0,40363384 |
| Lic1 | AX-111457218 | 4B | 533851398 | 0,33879781 | T | C | 27,5655289 | 4,1422E-07 | 0,00020298 | 12,3820727 | 0,56653471 | 0,37546902 |
| Lic1 | BobWhite_c30138_69 | 7D | 20933108 | 0,34224599 | T | C | 27,4724851 | 4,3193E-07 | 0,00020298 | 12,9299025 | 0,56311696 | 0,37018024 |
| Lic1 | TA003390-0807 | 2A | 787686566 | 0,46236559 | A | C | 26,6239101 | 6,3342E-07 | 0,00024513 | 12,5610084 | 0,58023713 | 0,39879258 |
| Lic1 | Kukri_c13045_302 | 5D | 244100832 | 0,44385027 | G | T | 26,1072585 | 8,0046E-07 | 0,00029809 | 12,3668218 | 0,57706298 | 0,39687195 |
| Lic1 | RAC875_c18659_402 | 6D | 32298303 | 0,12365591 | A | G | 25,3177141 | 1,1463E-06 | 0,00038788 | 12,03499 | 0,53006096 | 0,26105721 |
| Lic1 | JD_c1555_151 | 6B | 574415823 | 0,2513369 | G | T | 24,1704207 | 1,9374E-06 | 0,00057467 | 11,5553722 | 0,54722638 | 0,34772825 |
| Lic1 | IAAV4569 | 1B | 582731648 | 0,1657754 | G | A | 24,0914072 | 2,009E-06 | 0,00057467 | 11,5219499 | 0,53560634 | 0,30323655 |
| Lic1 | RFL_Contig3424_754 | UN | 32176331 | 0,39572193 | T | G | 23,957532 | 2,1365E-06 | 0,00058158 | 11,4652637 | 0,56684199 | 0,39056455 |
| Lic1 | AX-158542589 | 5A | 522662153 | 0,44565217 | G | A | 23,6639598 | 2,4455E-06 | 0,00064177 | 11,1438565 | 0,41610286 | 0,58825836 |
| Lic1 | AX-111584542 | UN | 222960893 | 0,12903226 | T | C | 23,6412298 | 2,4712E-06 | 0,00064177 | 11,2831595 | 0,52886754 | 0,2731244 |
| Lic2 |  |  |  |  |  |  |  |  |  |  |  |  |
| MCARI1 | BS00086534_51 | 2D | 656028039 | 0,33125 | G | A | 45,053206 | 2,2752E-10 | 4,4906E-06 | 15,2482105 | 0,63637179 | 0,31686555 |
| MCARI1 | Excalibur_c24638_380 | 5D | 359180839 | 0,12834225 | A | G | 41,0460425 | 1,1981E-09 | 8,4882E-06 | 18,1582664 | 0,57077052 | 0,11353305 |
| MCARI1 | AX-158551103 | 5A | 458638516 | 0,09625668 | T | C | 40,1942501 | 1,7127E-09 | 8,4882E-06 | 17,8487018 | 0,56157093 | 0,04749443 |
| MCARI1 | IAAV3697 | 4A | 38335364 | 0,17647059 | T | C | 39,180304 | 2,626E-09 | 8,6383E-06 | 17,4771393 | 0,58154115 | 0,18797124 |
| MCARI1 | wsnp_Ex_c36049_44083089 | 2A | 2925147 | 0,19672131 | C | T | 38,3430375 | 3,7435E-09 | 1,0555E-05 | 16,9323375 | 0,57968758 | 0,20447198 |
| MCARI1 | wsnp_Ex_c7216_12390889 | 7A | 524972333 | 0,34054054 | A | G | 37,3194254 | 5,7866E-09 | 1,4276E-05 | 16,7776005 | 0,61862492 | 0,30674836 |
| MCARI1 | TG0028 | 5D | 3622961 | 0,48648649 | G | A | 35,0235743 | 1,5499E-08 | 2,7809E-05 | 15,8012595 | 0,65392425 | 0,36699254 |
| MCARI1 | AX-95099091 | 5B | 424325426 | 0,13903743 | A | G | 34,5757184 | 1,8809E-08 | 2,8556E-05 | 15,7466038 | 0,56931841 | 0,15769707 |
| MCARI1 | AX-95168570 | 2B | 812643564 | 0,43646409 | G | C | 33,7365088 | 2,7066E-08 | 3,8157E-05 | 15,0634247 | 0,63538149 | 0,34991679 |
| MCARI1 | AX-109816649 | 5A | 521757324 | 0,34234234 | T | C | 33,0416461 | 3,663E-08 | 4,8197E-05 | 5,46097402 | 0,64514046 | 0,53669654 |
| MCARI1 | RAC875_c18659_402 | 6D | 32298303 | 0,12365591 | A | G | 32,6740616 | 4,3008E-08 | 5,3053E-05 | 15,0094061 | 0,56422732 | 0,14071782 |
| MCARI1 | AX-111584542 | UN | 222960893 | 0,12903226 | T | C | 30,7920211 | 9,8331E-08 | 0,00010782 | 14,1980064 | 0,56240772 | 0,15797231 |
| MCARI1 | IAAV149 | 1D | 429717285 | 0,14673913 | A | G | 29,9110387 | 1,4524E-07 | 0,00015087 | 13,7464637 | 0,56790602 | 0,18880452 |
| MCARI1 | TA003390-0807 | 2A | 787686566 | 0,46236559 | A | C | 29,2668539 | 1,9341E-07 | 0,00018178 | 13,6447155 | 0,63450003 | 0,36789783 |
| MCARI1 | Ra_c108731_554 | 5B | 61846046 | 0,17741935 | G | T | 28,8481114 | 2,3313E-07 | 0,00020915 | 13,426312 | 0,57150621 | 0,22638286 |
| MCARI1 | AX-158545574 | 1D | 455784529 | 0,38502674 | T | C | 28,6407975 | 2,5575E-07 | 0,00021947 | 13,4060525 | 0,61606294 | 0,34601596 |
| MCARI1 | Kukri_c2357_152 | 6A | 30503329 | 0,12972973 | G | A | 28,2676085 | 3,0224E-07 | 0,00024855 | 13,1768644 | 0,56040209 | 0,1706463 |
| MCARI1 | JD_c1555_151 | 6B | 574415823 | 0,2513369 | G | T | 27,8456261 | 3,6522E-07 | 0,00027724 | 13,0825456 | 0,58730073 | 0,28804862 |
| MCARI1 | AX-94964521 | 7A | 20912833 | 0,20540541 | A | G | 26,8684741 | 5,6713E-07 | 0,00038189 | 12,6747937 | 0,57821898 | 0,2602533 |
| MCARI1 | AX-158542764 | 5A | 621549770 | 0,49462366 | C | T | 26,6061705 | 6,3852E-07 | 0,00038189 | 12,5679757 | 0,63774525 | 0,38258648 |
| MCARI1 | BobWhite_c16397_524 | 5D | 498621954 | 0,49462366 | T | G | 26,6061705 | 6,3852E-07 | 0,00038189 | 12,5679757 | 0,63774525 | 0,38258648 |
| MCARI1 | RAC875_rep_c111494_195 | 1B | 665826070 | 0,42162162 | T | C | 25,7995968 | 9,2048E-07 | 0,00051907 | 12,2254464 | 0,40313776 | 0,65860862 |
| MCARI1 | AX-95208179 | 1D | 335597569 | 0,26344086 | C | T | 25,4770588 | 1,066E-06 | 0,00058442 | 12,0472879 | 0,58496481 | 0,30144705 |
| MCARI1 | AX-111457218 | 4B | 533851398 | 0,33879781 | T | C | 24,313882 | 1,814E-06 | 0,00071605 | 11,1843566 | 0,60464169 | 0,34858966 |
| MCARI1 | RAC875_c64377_732 | 2D | 573933688 | 0,40437158 | C | T | 23,8649115 | 2,2294E-06 | 0,00081082 | 11,3459193 | 0,40785234 | 0,65674495 |
| MCARI1 | IAAV4569 | 1B | 582731648 | 0,1657754 | G | A | 23,6558786 | 2,4546E-06 | 0,00081082 | 11,3372692 | 0,56595618 | 0,24100724 |
| MCARI | BS00086534_51 | 2D | 656028039 | 0,33125 | G | A | 43,2105405 | 4,86E-10 | 9,60E-06 | 14,7462684 | 0,07178078 | -0,0004235 |
| MCARI | AX-109816649 | 5A | 521757324 | 0,34234234 | T | C | 37,8599072 | 4,60E-09 | 3,39E-05 | 5,56891386 | 0,07372449 | 0,04123232 |
| MCARI | IAAV3697 | 4A | 38335364 | 0,17647059 | T | C | 36,3972941 | 8,58E-09 | 3,39E-05 | 16,4398098 | 0,05798452 | -0,0301883 |
| MCARI | Excalibur_c24638_380 | 5D | 359180839 | 0,12834225 | A | G | 35,4771544 | 1,27E-08 | 3,66E-05 | 16,0910796 | 0,05518507 | -0,04424021 |
| MCARI | AX-158551103 | 5A | 458638516 | 0,09625668 | T | C | 35,432466 | 1,30E-08 | 3,66E-05 | 16,0740687 | 0,05327181 | -0,0594186 |
| MCARI | AX-95168570 | 2B | 812643564 | 0,43646409 | G | C | 34,6023071 | 1,86E-08 | 4,59E-05 | 15,4814331 | 0,07176625 | 0,00491511 |
| MCARI | wsnp_Ex_c36049_44083089 | 2A | 2925147 | 0,19672131 | C | T | 32,7594074 | 4,14E-08 | 9,09E-05 | 14,6457552 | 0,05647028 | -0,02408036 |
| MCARI | TG0028 | 5D | 3622961 | 0,48648649 | G | A | 32,1014561 | 5,53E-08 | 9,41E-05 | 14,7291875 | 0,07388883 | 0,00989298 |
| MCARI | TA003390-0807 | 2A | 787686566 | 0,46236559 | A | C | 30,9462087 | 9,19E-08 | 0,00013947 | 14,3104338 | 0,07135664 | 0,00828922 |
| MCARI | RAC875_c18659_402 | 6D | 32298303 | 0,12365591 | A | G | 29,3604451 | 1,86E-07 | 0,00026154 | 13,6964877 | 0,05395304 | -0,03949841 |
| MCARI | Ra_c108731_554 | 5B | 61846046 | 0,17741936 | G | T | 28,8240535 | 2,36E-07 | 0,00031006 | 13,4486013 | 0,0562857 | -0,02350547 |
| MCARI | wsnp_Ex_c7216_12390889 | 7A | 524972333 | 0,34054054 | A | G | 28,3352192 | 2,93E-07 | 0,00033671 | 13,2695563 | 0,06440046 | 0,00033251 |
| MCARI | AX-95099091 | 5B | 424325426 | 0,13903743 | A | G | 27,9410234 | 3,50E-07 | 0,00036132 | 13,1214845 | 0,05449238 | -0,03230278 |
| MCARI | AX-111584542 | UN | 222960893 | 0,12903226 | T | C | 27,8108554 | 3,71E-07 | 0,00036132 | 12,9821667 | 0,05345593 | -0,03587266 |
| MCARI | AX-158545574 | 1D | 455784529 | 0,38502674 | T | C | 26,3998697 | 7,01E-07 | 0,00062891 | 12,4881201 | 0,06560538 | 0,00539976 |
| MCARI | BS00018740_51 | 5D | 542233108 | 0,42162162 | T | C | 25,4001387 | 1,10E-06 | 0,0009079 | 11,4614858 | 0,06755259 | 0,01045109 |
| MCARI | Kukri_c2357_152 | 6A | 30503329 | 0,12972973 | G | A | 24,9688304 | 1,34E-06 | 0,00102032 | 11,8640846 | 0,0531867 | -0,03224913 |
| NPCI |  |  |  |  |  |  |  |  |  |  |  |  |
| NPQI | IAAV3697 | 4A | 38335364 | 0,17647059 | T | C | 27,2832599 | 4,7035E-07 | 0,00464161 | 12,8522899 | -0,03106528 | -0,01106526 |
| NPQI | wsnp_Ex_c36049_44083089 | 2A | 2925147 | 0,19672131 | C | T | 25,9598091 | 8,5586E-07 | 0,00563072 | 12,3043886 | -0,03125963 | -0,01228899 |
| NPQI | AX-95168570 | 2B | 812643564 | 0,43646409 | G | C | 24,1931093 | 1,9174E-06 | 0,0094607 | 11,2251581 | -0,03368833 | -0,01909143 |
| OSAVI | wsnp_Ex_c36049_44083089 | 2A | 2925147 | 0,19672131 | C | T | 46,2574264 | 1,3897E-10 | 1,8332E-06 | 19,797389 | 0,5142988 | 0,24043193 |
| OSAVI | IAAV3697 | 4A | 38335364 | 0,17647059 | T | C | 44,5601357 | 2,7864E-10 | 1,8332E-06 | 19,4110949 | 0,51340146 | 0,23351372 |
| OSAVI | AX-158584871 | 5A | 535796991 | 0,2311828 | A | G | 42,3805748 | 6,8651E-10 | 2,8716E-06 | 18,2168601 | 0,52350202 | 0,27774761 |
| OSAVI | wsnp_Ex_c7216_12390889 | 7A | 524972333 | 0,34054054 | A | G | 41,8196057 | 8,6719E-10 | 2,8716E-06 | 18,4335502 | 0,53884732 | 0,31825318 |
| OSAVI | BS00086534_51 | 2D | 656028039 | 0,33125 | G | A | 41,8037095 | 8,7296E-10 | 2,8716E-06 | 14,6341151 | 0,54484542 | 0,33327698 |
| OSAVI | Excalibur_rep_c68688_103 | 5D | 45176107 | 0,14438503 | T | C | 37,4969736 | 5,3647E-09 | 1,5126E-05 | 16,8528017 | 0,50485015 | 0,2219909 |
| OSAVI | AX-158584345 | 5A | 524940962 | 0,25225225 | A | G | 35,7607399 | 1,1281E-08 | 2,474E-05 | 5,90381475 | 0,5140576 | 0,58931004 |
| OSAVI | Ra_c108731_554 | 5B | 61846046 | 0,17741935 | G | T | 33,1957723 | 3,4249E-08 | 6,1451E-05 | 15,1530678 | 0,50678145 | 0,25936491 |
| OSAVI | RAC875_rep_c111494_195 | 1B | 665826070 | 0,42162162 | T | C | 31,9883469 | 5,8073E-08 | 7,6412E-05 | 14,7113688 | 0,38313104 | 0,57223288 |
| OSAVI | AX-95168570 | 2B | 812643564 | 0,43646409 | G | C | 31,4260411 | 7,4351E-08 | 8,3599E-05 | 14,2327343 | 0,54359984 | 0,35639703 |
| OSAVI | AX-95208179 | 1D | 335597569 | 0,26344086 | C | T | 31,3119516 | 7,8181E-08 | 8,3599E-05 | 14,4176256 | 0,518023 | 0,30872353 |
| OSAVI | IAAV149 | 1D | 429717285 | 0,14673913 | A | G | 30,9946905 | 8,9916E-08 | 8,4508E-05 | 14,2017716 | 0,50179118 | 0,24177505 |
| OSAVI | AX-158545574 | 1D | 455784529 | 0,38502674 | T | C | 30,55162 | 1,0935E-07 | 9,384E-05 | 14,1736907 | 0,53615238 | 0,34878132 |
| OSAVI | AX-95099091 | 5B | 424325426 | 0,13903743 | A | G | 30,0387287 | 1,3724E-07 | 0,00010774 | 13,9689854 | 0,50038348 | 0,23877066 |
| OSAVI | IAAV912 | 7A | 20370858 | 0,28877005 | G | T | 29,9631832 | 1,4192E-07 | 0,00010774 | 13,9387512 | 0,52162212 | 0,32211177 |
| OSAVI | AX-158542802 | 5A | 458637478 | 0,14606742 | A | C | 29,9631086 | 1,4192E-07 | 0,00010774 | 13,682724 | 0,49857616 | 0,23877066 |
| OSAVI | RAC875_c64377_732 | 2D | 573933688 | 0,40437158 | C | T | 29,8322261 | 1,5041E-07 | 0,00010995 | 13,7503861 | 0,38611947 | 0,57098241 |
| OSAVI | RAC875_c33083_451 | 3B | 841569821 | 0,26344086 | A | G | 28,9524432 | 2,2252E-07 | 0,00015143 | 13,478206 | 0,51619613 | 0,3138313 |
| OSAVI | Kukri_rep_c102102_273 | 1A | 571167863 | 0,40322581 | C | T | 28,876687 | 2,3017E-07 | 0,00015143 | 13,4058426 | 0,38943796 | 0,57065752 |
| OSAVI | AX-158569026 | 1A | 586999805 | 0,32795699 | T | C | 28,577942 | 2,6304E-07 | 0,00016747 | 13,2713011 | 0,52418705 | 0,33578881 |
| OSAVI | BobWhite_c16397_524 | 5D | 498621954 | 0,49462366 | T | G | 27,3946984 | 4,4732E-07 | 0,00025225 | 12,8869291 | 0,54972497 | 0,37537537 |
| OSAVI | AX-111457218 | 4B | 533851398 | 0,33879781 | T | C | 26,8776626 | 5,6478E-07 | 0,0002576 | 12,1006655 | 0,52958575 | 0,34993394 |
| OSAVI | BobWhite_c30138_69 | 7D | 20933108 | 0,34224599 | T | C | 26,7198398 | 6,0653E-07 | 0,0002576 | 12,620376 | 0,52606899 | 0,34473893 |
| OSAVI | TA003390-0807 | 2A | 787686566 | 0,46236559 | A | C | 26,4796624 | 6,7614E-07 | 0,00026424 | 12,5032847 | 0,5429617 | 0,3707505 |
| OSAVI | Kukri_c13045_302 | 5D | 244100832 | 0,44385027 | G | T | 25,7862546 | 9,2608E-07 | 0,00034487 | 12,2333663 | 0,53968059 | 0,36919273 |
| OSAVI | RAC875_c18659_402 | 6D | 32298303 | 0,12365591 | A | G | 25,3164539 | 1,1469E-06 | 0,00039665 | 12,0348374 | 0,49539776 | 0,23949659 |
| OSAVI | JD_c1555_151 | 6B | 574415823 | 0,2513369 | G | T | 24,9095944 | 1,381E-06 | 0,00045359 | 11,8668203 | 0,51234729 | 0,32002464 |
| OSAVI | IAAV4569 | 1B | 582731648 | 0,1657754 | G | A | 24,1699503 | 1,9378E-06 | 0,00055431 | 11,5551733 | 0,50070742 | 0,27933614 |
| OSAVI | AX-111584542 | UN | 222960893 | 0,12903226 | T | C | 23,6718845 | 2,4366E-06 | 0,00067829 | 11,2945138 | 0,49424036 | 0,25083096 |
| OSAVI | AX-158542589 | 5A | 522662153 | 0,44565217 | G | A | 23,6236748 | 2,4913E-06 | 0,00068242 | 11,1295041 | 0,38705764 | 0,5507271 |
| PRI |  |  |  |  |  |  |  |  |  |  |  |  |
| RDVI | wsnp_Ex_c36049_44083089 | 2A | 2925147 | 0,19672131 | C | T | 44,03756 | 3,46E-10 | 2,86E-06 | 19,007619 | 0,44613859 | 0,20391819 |
| RDVI | IAAV3697 | 4A | 38335364 | 0,17647059 | T | C | 43,4804384 | 4,35E-10 | 2,86E-06 | 19,0302674 | 0,44604568 | 0,19587908 |
| RDVI | AX-158584871 | 5A | 535796991 | 0,2311828 | A | G | 41,7049133 | 9,10E-10 | 4,49E-06 | 17,9883586 | 0,45525574 | 0,23480605 |
| RDVI | BS00086534_51 | 2D | 656028039 | 0,33125 | G | A | 41,0588514 | 1,19E-09 | 4,70E-06 | 14,3739928 | 0,47457323 | 0,2853834 |
| RDVI | wsnp_Ex_c7216_12390889 | 7A | 524972333 | 0,34054054 | A | G | 39,2761652 | 2,52E-09 | 7,72E-06 | 17,5088668 | 0,46788629 | 0,27381208 |
| RDVI | Excalibur_rep_c68688_103 | 5D | 45176107 | 0,14438503 | T | C | 39,0802691 | 2,74E-09 | 7,72E-06 | 17,4402991 | 0,43940296 | 0,17965073 |
| RDVI | AX-158584345 | 5A | 524940962 | 0,25225225 | A | G | 33,719038 | 2,73E-08 | 5,38E-05 | 5,57266208 | 0,44720886 | 0,5100727 |
| RDVI | Ra_c108731_554 | 5B | 61846046 | 0,17741936 | G | T | 32,8510598 | 3,98E-08 | 7,14E-05 | 15,0203236 | 0,44034279 | 0,21797787 |
| RDVI | AX-95168570 | 2B | 812643564 | 0,43646409 | G | C | 31,4260639 | 7,44E-08 | 9,17E-05 | 14,2294797 | 0,47382573 | 0,30485141 |
| RDVI | AX-95099091 | 5B | 424325426 | 0,13903743 | A | G | 30,6871725 | 1,03E-07 | 0,00011209 | 14,2276298 | 0,43503635 | 0,19669968 |
| RDVI | AX-158542802 | 5A | 458637478 | 0,14606742 | A | C | 30,6368202 | 1,05E-07 | 0,00011209 | 13,9560969 | 0,43357625 | 0,19669968 |
| RDVI | RAC875_rep_c111494_195 | 1B | 665826070 | 0,42162162 | T | C | 30,1170201 | 1,33E-07 | 0,00011824 | 13,968233 | 0,33068099 | 0,49701565 |
| RDVI | AX-95208179 | 1D | 335597569 | 0,26344086 | C | T | 30,0558763 | 1,36E-07 | 0,00011824 | 13,9209111 | 0,44979959 | 0,2641464 |
| RDVI | IAAV149 | 1D | 429717285 | 0,14673913 | A | G | 30,0350133 | 1,37E-07 | 0,00011824 | 13,8302359 | 0,43557582 | 0,20394718 |
| RDVI | AX-158545574 | 1D | 455784529 | 0,38502674 | T | C | 30,0297643 | 1,38E-07 | 0,00011824 | 13,9653989 | 0,46654239 | 0,29864818 |
| RDVI | IAAV912 | 7A | 20370858 | 0,28877005 | G | T | 29,0842939 | 2,10E-07 | 0,00016564 | 13,5854403 | 0,45324273 | 0,27544002 |
| RDVI | RAC875_c64377_732 | 2D | 573933688 | 0,40437159 | C | T | 28,0148416 | 3,39E-07 | 0,00023861 | 13,0194847 | 0,33337684 | 0,49575095 |
| RDVI | RAC875_c33083_451 | 3B | 841569821 | 0,26344086 | A | G | 27,3956807 | 4,47E-07 | 0,00030246 | 12,847718 | 0,44787625 | 0,2695239 |
| RDVI | BobWhite_c16397_524 | 5D | 498621954 | 0,49462366 | T | G | 27,2611191 | 4,75E-07 | 0,00030246 | 12,8316289 | 0,47909627 | 0,32204743 |
| RDVI | Kukri_rep_c102102_273 | 1A | 571167863 | 0,40322581 | C | T | 27,0464821 | 5,23E-07 | 0,00032279 | 12,6627497 | 0,33642372 | 0,495412 |
| RDVI | JD_c1555_151 | 6B | 574415823 | 0,2513369 | G | T | 26,6718468 | 6,20E-07 | 0,00037021 | 12,6005641 | 0,44686238 | 0,26796406 |
| RDVI | TA003390-0807 | 2A | 787686566 | 0,46236559 | A | C | 26,4397154 | 6,88E-07 | 0,00038825 | 12,4892219 | 0,47317246 | 0,31780257 |
| RDVI | RAC875_c18659_402 | 6D | 32298303 | 0,12365591 | A | G | 25,5409573 | 1,04E-06 | 0,00041015 | 12,1291488 | 0,43037101 | 0,19846299 |
| RDVI | AX-158569026 | 1A | 586999805 | 0,32795699 | T | C | 25,540163 | 1,04E-06 | 0,00041015 | 12,039007 | 0,45362318 | 0,29164299 |
| RDVI | AX-111457218 | 4B | 533851398 | 0,33879781 | T | C | 25,3643451 | 1,12E-06 | 0,00041015 | 11,4838575 | 0,45995017 | 0,3020132 |
| RDVI | BobWhite_c30138_69 | 7D | 20933108 | 0,34224599 | T | C | 25,2251186 | 1,20E-06 | 0,00042142 | 11,9990983 | 0,45652399 | 0,29691551 |
| RDVI | Kukri_c13045_302 | 5D | 244100832 | 0,44385027 | G | T | 25,0947342 | 1,27E-06 | 0,00043941 | 11,9444851 | 0,46939629 | 0,31732325 |
| RDVI | AX-111584542 | UN | 222960893 | 0,12903226 | T | C | 23,9329957 | 2,16E-06 | 0,00068636 | 11,4023377 | 0,4292971 | 0,20852371 |
| SIPI | wsnp_Ex_c36049_44083089 | 2A | 2925147 | 0,19672131 | C | T | 48,3056761 | 6,05E-11 | 1,19E-06 | 20,5077095 | 0,54609574 | 0,26150655 |
| SIPI | IAAV3697 | 4A | 38335364 | 0,17647059 | T | C | 44,3420384 | 3,05E-10 | 2,01E-06 | 19,3344573 | 0,54397824 | 0,25879353 |
| SIPI | AX-158584871 | 5A | 535796991 | 0,2311828 | A | G | 40,900924 | 1,27E-09 | 4,30E-06 | 17,7525523 | 0,55345249 | 0,30575985 |
| SIPI | BS00086534_51 | 2D | 656028039 | 0,33125 | G | A | 40,8896078 | 1,28E-09 | 4,30E-06 | 13,9278943 | 0,5763566 | 0,36624553 |
| SIPI | wsnp_Ex_c7216_12390889 | 7A | 524972333 | 0,34054054 | A | G | 40,8384466 | 1,31E-09 | 4,30E-06 | 18,0732616 | 0,56900352 | 0,34600397 |
| SIPI | AX-158584345 | 5A | 524940962 | 0,25225225 | A | G | 35,3120017 | 1,37E-08 | 3,86E-05 | 5,70099947 | 0,54324451 | 0,62008471 |
| SIPI | Excalibur_c24638_380 | 5D | 359180839 | 0,12834225 | A | G | 33,3429066 | 3,21E-08 | 6,34E-05 | 15,2708907 | 0,53072643 | 0,24185117 |
| SIPI | IAAV149 | 1D | 429717285 | 0,14673913 | A | G | 32,3636566 | 4,93E-08 | 8,10E-05 | 14,7317101 | 0,53268164 | 0,26231646 |
| SIPI | Ra_c108731_554 | 5B | 61846046 | 0,17741936 | G | T | 31,7339934 | 6,49E-08 | 9,32E-05 | 14,5967843 | 0,5366285 | 0,28870681 |
| SIPI | RAC875_rep_c111494_195 | 1B | 665826070 | 0,42162162 | T | C | 31,4705166 | 7,29E-08 | 9,51E-05 | 14,5149762 | 0,41176813 | 0,6035405 |
| SIPI | AX-95168570 | 2B | 812643564 | 0,43646409 | G | C | 31,3435524 | 7,71E-08 | 9,51E-05 | 14,1921872 | 0,57507597 | 0,38421311 |
| SIPI | AX-95208179 | 1D | 335597569 | 0,26344086 | C | T | 30,9253316 | 9,27E-08 | 9,79E-05 | 14,2780949 | 0,54866302 | 0,33601328 |
| SIPI | TG0028 | 5D | 3622961 | 0,48648649 | G | A | 30,7306905 | 1,01E-07 | 9,79E-05 | 14,1935095 | 0,58566085 | 0,39829682 |
| SIPI | RAC875_c64377_732 | 2D | 573933688 | 0,40437159 | C | T | 29,713195 | 1,59E-07 | 0,00013608 | 13,7181019 | 0,41439962 | 0,6029248 |
| SIPI | AX-158545574 | 1D | 455784529 | 0,38502674 | T | C | 29,5150243 | 1,73E-07 | 0,00014242 | 13,7589544 | 0,56621967 | 0,37774408 |
| SIPI | IAAV912 | 7A | 20370858 | 0,28877005 | G | T | 29,2438809 | 1,95E-07 | 0,00015427 | 13,6498092 | 0,55185787 | 0,35029146 |
| SIPI | Kukri_rep_c102102_273 | 1A | 571167863 | 0,40322581 | C | T | 28,9019909 | 2,28E-07 | 0,00016637 | 13,4256076 | 0,41754096 | 0,60269476 |
| SIPI | AX-95099091 | 5B | 424325426 | 0,13903743 | A | G | 28,7484811 | 2,44E-07 | 0,00017181 | 13,4496774 | 0,53009046 | 0,26801045 |
| SIPI | AX-158542802 | 5A | 458637478 | 0,14606742 | A | C | 28,4957453 | 2,73E-07 | 0,00018573 | 13,0967571 | 0,52743003 | 0,26801045 |
| SIPI | AX-158569026 | 1A | 586999805 | 0,32795699 | T | C | 27,9677305 | 3,46E-07 | 0,00022746 | 13,0271394 | 0,55452179 | 0,36395552 |
| SIPI | AX-111457218 | 4B | 533851398 | 0,33879781 | T | C | 27,7781757 | 3,76E-07 | 0,00023583 | 12,4550169 | 0,561307 | 0,37519503 |
| SIPI | RAC875_c33083_451 | 3B | 841569821 | 0,26344086 | A | G | 27,4613478 | 4,34E-07 | 0,00025963 | 12,885504 | 0,54586067 | 0,34384842 |
| SIPI | BobWhite_c16397_524 | 5D | 498621954 | 0,49462366 | T | G | 27,0669418 | 5,19E-07 | 0,00028428 | 12,7541217 | 0,58075041 | 0,40366857 |
| SIPI | TA003390-0807 | 2A | 787686566 | 0,46236559 | A | C | 26,4726586 | 6,78E-07 | 0,0003342 | 12,4949381 | 0,57413141 | 0,39837399 |
| SIPI | BobWhite_c30138_69 | 7D | 20933108 | 0,34224599 | T | C | 26,2757441 | 7,42E-07 | 0,00034038 | 12,4367065 | 0,55654797 | 0,37277242 |
| SIPI | Kukri_c13045_302 | 5D | 244100832 | 0,44385027 | G | T | 25,2185987 | 1,20E-06 | 0,00043036 | 11,9963689 | 0,57015537 | 0,39779129 |
| SIPI | IAAV4569 | 1B | 582731648 | 0,1657754 | G | A | 24,6949367 | 1,52E-06 | 0,00048319 | 11,7766014 | 0,53147527 | 0,30331269 |
| SIPI | tplb0032i02_1388 | 2A | 7599387 | 0,1827957 | C | T | 23,5925622 | 2,53E-06 | 0,00070254 | 11,2382143 | 0,53150046 | 0,31650248 |
| SPRI |  |  |  |  |  |  |  |  |  |  |  |  |
| SR | wsnp_Ex_c36049_44083089 | 2A | 2925147 | 0,19672131 | C | T | 39,3656849 | 2,4283E-09 | 4,7926E-05 | 17,4917695 | 3,9613245 | 2,47475523 |
| SR | wsnp_Ex_c7216_12390889 | 7A | 524972333 | 0,34054054 | A | G | 34,8333948 | 1,6825E-08 | 0,00011345 | 15,8305427 | 4,08010227 | 2,9000292 |
| SR | IAAV3697 | 4A | 38335364 | 0,17647059 | T | C | 34,1121791 | 2,2992E-08 | 0,00011345 | 15,5683629 | 3,93516876 | 2,48822378 |
| SR | AX-158584345 | 5A | 524940962 | 0,25225225 | A | G | 31,9909051 | 5,8008E-08 | 0,00022898 | 6,09574936 | 3,91191041 | 4,47731259 |
| SR | BS00086534_51 | 2D | 656028039 | 0,33125 | G | A | 30,3084671 | 1,2178E-07 | 0,00034318 | 11,4682348 | 4,08929343 | 3,00727552 |
| SR | AX-158584871 | 5A | 535796991 | 0,2311828 | A | G | 30,0182834 | 1,3849E-07 | 0,00034318 | 13,6948173 | 3,97836199 | 2,74838514 |
| SR | IAAV149 | 1D | 429717285 | 0,14673913 | A | G | 30,0083622 | 1,391E-07 | 0,00034318 | 13,8628526 | 3,89619581 | 2,41323605 |
| SR | AX-158569026 | 1A | 586999805 | 0,32795699 | T | C | 28,3280005 | 2,9417E-07 | 0,00062434 | 13,1316083 | 4,023759 | 2,94201668 |
| SR | AX-95208179 | 1D | 335597569 | 0,26344086 | C | T | 28,0204137 | 3,3766E-07 | 0,00062434 | 13,1308664 | 3,97929422 | 2,82622238 |
| SR | RAC875_c33083_451 | 3B | 841569821 | 0,26344086 | A | G | 27,2521239 | 4,7699E-07 | 0,00065774 | 12,8170587 | 3,97564223 | 2,83643304 |
| SR | RAC875_rep_c111494_195 | 1B | 665826070 | 0,42162162 | T | C | 26,911437 | 5,5623E-07 | 0,00065774 | 12,5535105 | 3,23956727 | 4,24754752 |
| SR | BobWhite_c30138_69 | 7D | 20933108 | 0,34224599 | T | C | 26,909916 | 5,5661E-07 | 0,00065774 | 12,6987526 | 4,03918097 | 2,98918928 |
| SR | Ra_c108731_554 | 5B | 61846046 | 0,17741935 | G | T | 26,442095 | 6,8774E-07 | 0,00065774 | 12,483733 | 3,90553323 | 2,60914003 |
| SR | RAC875_c64377_732 | 2D | 573933688 | 0,40437158 | C | T | 26,2254529 | 7,5868E-07 | 0,00065774 | 12,1576007 | 3,24555519 | 4,24785009 |
| SR | Excalibur_c24638_380 | 5D | 359180839 | 0,12834225 | A | G | 25,7345201 | 9,4812E-07 | 0,0007259 | 12,2118199 | 3,86728538 | 2,40666066 |
| SR | AX-108742004 | 7A | 21131825 | 0,3368984 | T | C | 25,3212043 | 1,1444E-06 | 0,0007259 | 12,0393017 | 4,02557884 | 2,9992952 |
| SR | Kukri_c16352_687 | 3D | 611641089 | 0,17112299 | G | A | 24,8844672 | 1,3969E-06 | 0,0007259 | 11,8562691 | 3,898546 | 2,62039825 |
| SR | Kukri_c13045_302 | 5D | 244100832 | 0,44385027 | G | T | 24,7562074 | 1,4813E-06 | 0,0007259 | 11,8023718 | 4,10888084 | 3,14221404 |
| SR | IAAV4569 | 1B | 582731648 | 0,1657754 | G | A | 24,5331175 | 1,6406E-06 | 0,0007259 | 11,7084678 | 3,89306856 | 2,60673156 |
| SR | Kukri_rep_c102102_273 | 1A | 571167863 | 0,40322581 | C | T | 24,4478407 | 1,7059E-06 | 0,0007259 | 11,5856445 | 3,27883319 | 4,25130386 |
| SR | AX-158542802 | 5A | 458637478 | 0,14606742 | A | C | 24,4322335 | 1,7182E-06 | 0,0007259 | 11,2814371 | 3,86610716 | 2,50384855 |
| SR | AX-158545574 | 1D | 455784529 | 0,38502674 | T | C | 24,4190477 | 1,7286E-06 | 0,0007259 | 11,6603757 | 4,05755412 | 3,07650902 |
| SR | AX-95099091 | 5B | 424325426 | 0,13903743 | A | G | 23,8672839 | 2,227E-06 | 0,00081035 | 11,4270093 | 3,86973485 | 2,50384855 |
| TCARI | RAC875_rep_c102342_470 | 5B | 576702400 | 0,09090909 | G | A | 45,2463484 | 2,1018E-10 | 4,1484E-06 | 19,6512773 | -0,15781401 | -0,29958061 |
| TCARI | AX-95077341 | 6B | 702874103 | 0,07526882 | G | C | 40,2689169 | 1,6598E-09 | 1,638E-05 | 17,8758708 | -0,15956999 | -0,30729991 |
| TCARI | AX-94406390 | 7B | 477131416 | 0,0802139 | G | A | 38,164766 | 4,0378E-09 | 2,6565E-05 | 17,1016091 | -0,15947431 | -0,29944467 |
| TCARI | AX-94601987 | 2A | 199424810 | 0,06989247 | A | C | 37,1062392 | 6,3378E-09 | 3,1272E-05 | 16,6960044 | -0,16054626 | -0,3082768 |
| TCARI | BS00027770_51 | 6D | 475964186 | 0,12299465 | A | G | 31,1894238 | 8,2518E-08 | 0,00027144 | 14,4268962 | -0,1576247 | -0,2639479 |
| TCARI | wsnp_Ex_c36049_44083089 | 2A | 2925147 | 0,19672131 | C | T | 30,7161191 | 1,0168E-07 | 0,0002867 | 14,214769 | -0,15357491 | -0,24171717 |
| TCARI | Kukri_c73802_205 | 6D | 4572453 | 0,12941176 | G | A | 30,218131 | 1,2675E-07 | 0,00029548 | 11,7940351 | -0,15522074 | -0,25353311 |
| TCARI | wsnp_RFL_Contig429_4978628 | 3B | 547032409 | 0,14438503 | A | G | 30,0802539 | 1,3474E-07 | 0,00029548 | 13,9855953 | -0,15657819 | -0,25439783 |
| TCARI | Kukri_c5757_530 | 7A | 94404713 | 0,07526882 | A | G | 24,2250187 | 1,8895E-06 | 0,00372926 | 11,5705277 | -0,1619329 | -0,28078437 |
| ZMI | AX-158584345 | 5A | 524940962 | 0,25225225 | A | G | 41,1965847 | 1,1249E-09 | 1,6347E-05 | 7,91403742 | 2,00807659 | 2,24405171 |
| ZMI | wsnp_Ex_c36049_44083089 | 2A | 2925147 | 0,19672131 | C | T | 40,2736899 | 1,6565E-09 | 1,6347E-05 | 17,8199722 | 2,03150359 | 1,5499215 |
| ZMI | wsnp_Ex_c7216_12390889 | 7A | 524972333 | 0,34054054 | A | G | 38,6453275 | 3,2931E-09 | 2,1666E-05 | 17,2559042 | 2,07314456 | 1,67773299 |
| ZMI | IAAV3697 | 4A | 38335364 | 0,17647059 | T | C | 33,9481116 | 2,4689E-08 | 9,7457E-05 | 15,5050945 | 2,02156969 | 1,55812716 |
| ZMI | AX-158584871 | 5A | 535796991 | 0,2311828 | A | G | 33,0699737 | 3,618E-08 | 0,00010359 | 14,9061755 | 2,0393109 | 1,62744209 |
| ZMI | RAC875_rep_c111494_195 | 1B | 665826070 | 0,42162162 | T | C | 33,0348701 | 3,6738E-08 | 0,00010359 | 14,9304526 | 1,78608342 | 2,13891197 |
| ZMI | AX-95208179 | 1D | 335597569 | 0,26344086 | C | T | 32,5858796 | 4,4699E-08 | 0,00011028 | 14,9164141 | 2,0416085 | 1,64719263 |
| ZMI | RAC875_c33083_451 | 3B | 841569821 | 0,26344086 | A | G | 32,0191654 | 5,7293E-08 | 0,0001186 | 14,6952713 | 2,04083529 | 1,64935446 |
| ZMI | RAC875_c64377_732 | 2D | 573933688 | 0,40437158 | C | T | 31,1578458 | 8,3674E-08 | 0,0001311 | 13,9994205 | 1,79017711 | 2,13535839 |
| ZMI | AX-158569026 | 1A | 586999805 | 0,32795699 | T | C | 31,0863702 | 8,6353E-08 | 0,0001311 | 14,1243664 | 2,0534073 | 1,69338797 |
| ZMI | BobWhite_c16397_524 | 5D | 498621954 | 0,49462366 | T | G | 29,341421 | 1,871E-07 | 0,00024618 | 13,6743059 | 2,10327927 | 1,77054521 |
| ZMI | IAAV149 | 1D | 429717285 | 0,14673913 | A | G | 29,1866059 | 2,0045E-07 | 0,00024727 | 13,5528331 | 2,00945104 | 1,53885905 |
| ZMI | Kukri_rep_c102102_273 | 1A | 571167863 | 0,40322581 | C | T | 28,8587069 | 2,3203E-07 | 0,00026938 | 13,320415 | 1,80110968 | 2,13574015 |
| ZMI | BS00086534_51 | 2D | 656028039 | 0,33125 | G | A | 28,7285742 | 2,4591E-07 | 0,00026964 | 11,0041429 | 2,06585487 | 1,72513629 |
| ZMI | AX-158545574 | 1D | 455784529 | 0,38502674 | T | C | 28,2701023 | 3,019E-07 | 0,00031361 | 13,2555393 | 2,06904163 | 1,7333353 |
| ZMI | Excalibur_rep_c68688_103 | 5D | 45176107 | 0,14438503 | T | C | 26,6054151 | 6,3874E-07 | 0,00052117 | 12,5731258 | 2,00514058 | 1,55249763 |
| ZMI | BobWhite_c30138_69 | 7D | 20933108 | 0,34224599 | T | C | 26,2383951 | 7,5424E-07 | 0,00053166 | 12,4212244 | 2,05385115 | 1,72056622 |
| ZMI | AX-158542802 | 5A | 458637478 | 0,14606742 | A | C | 25,3201365 | 1,145E-06 | 0,0006219 | 11,4803468 | 1,99853112 | 1,55762324 |
| ZMI | IAAV4569 | 1B | 582731648 | 0,1657754 | G | A | 25,2220775 | 1,1974E-06 | 0,0006219 | 11,9978253 | 2,00906513 | 1,59115382 |
| ZMI | IAAV912 | 7A | 20370858 | 0,28877005 | G | T | 24,9940457 | 1,3287E-06 | 0,00063239 | 11,902264 | 2,03841826 | 1,69685741 |
| ZMI | TA011876-1039 | 3D | 612524357 | 0,17204301 | A | G | 24,7699909 | 1,472E-06 | 0,00063239 | 11,808034 | 2,01036606 | 1,60076129 |
| ZMI | AX-95099091 | 5B | 424325426 | 0,13903743 | A | G | 24,5506991 | 1,6274E-06 | 0,00066917 | 11,7158755 | 2,0015014 | 1,55762324 |
| ZMI | Kukri_c13045_302 | 5D | 244100832 | 0,44385027 | G | T | 24,3388161 | 1,7934E-06 | 0,00070791 | 11,6265185 | 2,07645837 | 1,76853324 |
| ZMI | Ra_c108731_554 | 5B | 61846046 | 0,17741935 | G | T | 24,2473266 | 1,8702E-06 | 0,00072378 | 11,5416961 | 2,00867809 | 1,60863803 |
