## Supplementary Table S6 for "Wheat diversity reveals new genomic loci and candidate genes for vegetation indices using genome-wide association analysis"

**Table S6** Candidate genes and their functional annotations localizing at the 2A QTL peak

| **Transcript stable ID** | **Gene description** | **Gene start (bp)** | **Gene end (bp)** | **Interpro ID** | **Interpro Description** |
| --- | --- | --- | --- | --- | --- |
| TraesCS2A02G004600.1 |  | 2463117 | 2466740 | [IPR013087](http://www.ebi.ac.uk/interpro/entry/IPR013087) | Zinc finger C2H2-type |
| TraesCS2A02G004700.1 |  | 2476815 | 2481845 | [IPR000467](http://www.ebi.ac.uk/interpro/entry/IPR000467) | G-patch domain |
|  |  |  |  | [IPR013087](http://www.ebi.ac.uk/interpro/entry/IPR013087) | Zinc finger C2H2-type |
| TraesCS2A02G004800.1 |  | 2481972 | 2487182 | [IPR017423](http://www.ebi.ac.uk/interpro/entry/IPR017423) | tRNA (adenine(58)-N(1))-methyltransferase non-catalytic subunit TRM6 |
| TraesCS2A02G004800.2 |  | 2481972 | 2487182 | [IPR017423](http://www.ebi.ac.uk/interpro/entry/IPR017423) | tRNA (adenine(58)-N(1))-methyltransferase non-catalytic subunit TRM6 |
| TraesCS2A02G004900.1 |  | 2488830 | 2495245 | [IPR002182](http://www.ebi.ac.uk/interpro/entry/IPR002182) | NB-ARC |
|  |  |  |  | [IPR003593](http://www.ebi.ac.uk/interpro/entry/IPR003593) | AAA+ ATPase domain |
|  |  |  |  | [IPR027417](http://www.ebi.ac.uk/interpro/entry/IPR027417) | P-loop containing nucleoside triphosphate hydrolase |
|  |  |  |  | [IPR032675](http://www.ebi.ac.uk/interpro/entry/IPR032675) | Leucine-rich repeat domain superfamily |
|  |  |  |  | [IPR036388](http://www.ebi.ac.uk/interpro/entry/IPR036388) | Winged helix-like DNA-binding domain superfamily |
|  |  |  |  | [IPR041118](http://www.ebi.ac.uk/interpro/entry/IPR041118) | Disease resistance, N-terminal |
|  |  |  |  | [IPR042197](http://www.ebi.ac.uk/interpro/entry/IPR042197) | Apoptotic protease-activating factors, helical domain |
|  |  |  |  | [IPR055414](http://www.ebi.ac.uk/interpro/entry/IPR055414) | Disease resistance R13L4/SHOC-2-like, LRR domain |
| TraesCS2A02G005000.1 |  | 2637762 | 2653834 | [IPR001214](http://www.ebi.ac.uk/interpro/entry/IPR001214) | SET domain |
|  |  |  |  | [IPR003616](http://www.ebi.ac.uk/interpro/entry/IPR003616) | Post-SET domain |
|  |  |  |  | [IPR006560](http://www.ebi.ac.uk/interpro/entry/IPR006560) | AWS domain |
|  |  |  |  | [IPR011124](http://www.ebi.ac.uk/interpro/entry/IPR011124) | Zinc finger, CW-type |
|  |  |  |  | [IPR044437](http://www.ebi.ac.uk/interpro/entry/IPR044437) | SETD2/Set2, SET domain |
|  |  |  |  | [IPR046341](http://www.ebi.ac.uk/interpro/entry/IPR046341) | SET domain superfamily |
|  |  |  |  | [IPR050777](http://www.ebi.ac.uk/interpro/entry/IPR050777) | SET2 Histone-Lysine N-Methyltransferase |
| TraesCS2A02G005100.1 |  | 2655708 | 2657588 | [IPR001128](http://www.ebi.ac.uk/interpro/entry/IPR001128) | Cytochrome P450 |
|  |  |  |  | [IPR002401](http://www.ebi.ac.uk/interpro/entry/IPR002401) | Cytochrome P450, E-class, group I |
|  |  |  |  | [IPR017972](http://www.ebi.ac.uk/interpro/entry/IPR017972) | Cytochrome P450, conserved site |
|  |  |  |  | [IPR036396](http://www.ebi.ac.uk/interpro/entry/IPR036396) | Cytochrome P450 superfamily |
| TraesCS2A02G005200.1 | Phospho-transferase | 2657374 | 2660973 | [IPR001312](http://www.ebi.ac.uk/interpro/entry/IPR001312) | Hexokinase |
|  |  |  |  | [IPR019807](http://www.ebi.ac.uk/interpro/entry/IPR019807) | Hexokinase, binding site |
|  |  |  |  | [IPR022672](http://www.ebi.ac.uk/interpro/entry/IPR022672) | Hexokinase, N-terminal |
|  |  |  |  | [IPR022673](http://www.ebi.ac.uk/interpro/entry/IPR022673) | Hexokinase, C-terminal |
|  |  |  |  | [IPR043129](http://www.ebi.ac.uk/interpro/entry/IPR043129) | ATPase, nucleotide binding domain |
| TraesCS2A02G005300.1 |  | 2693455 | 2697385 |  |  |
| TraesCS2A02G005400.1 |  | 2700528 | 2700923 |  |  |
| TraesCS2A02G005500.1 |  | 2708175 | 2708888 | [IPR004864](http://www.ebi.ac.uk/interpro/entry/IPR004864) | Late embryogenesis abundant protein, LEA_2 subgroup |
|  |  |  |  | [IPR044839](http://www.ebi.ac.uk/interpro/entry/IPR044839) | Protein NDR1-like |
| TraesCS2A02G005600.1 |  | 2713730 | 2714453 |  |  |
| TraesCS2A02G005700.1 |  | 2726159 | 2727824 | [IPR006094](http://www.ebi.ac.uk/interpro/entry/IPR006094) | FAD linked oxidase, N-terminal |
|  |  |  |  | [IPR012951](http://www.ebi.ac.uk/interpro/entry/IPR012951) | Berberine/berberine-like |
|  |  |  |  | [IPR016166](http://www.ebi.ac.uk/interpro/entry/IPR016166) | FAD-binding domain, PCMH-type |
|  |  |  |  | [IPR016169](http://www.ebi.ac.uk/interpro/entry/IPR016169) | FAD-binding, type PCMH, subdomain 2 |
|  |  |  |  | [IPR036318](http://www.ebi.ac.uk/interpro/entry/IPR036318) | FAD-binding, type PCMH-like superfamily |
| TraesCS2A02G005800.1 |  | 2739129 | 2740683 | [IPR001128](http://www.ebi.ac.uk/interpro/entry/IPR001128) | Cytochrome P450 |
|  |  |  |  | [IPR002401](http://www.ebi.ac.uk/interpro/entry/IPR002401) | Cytochrome P450, E-class, group I |
|  |  |  |  | [IPR017972](http://www.ebi.ac.uk/interpro/entry/IPR017972) | Cytochrome P450, conserved site |
|  |  |  |  | [IPR036396](http://www.ebi.ac.uk/interpro/entry/IPR036396) | Cytochrome P450 superfamily |
| TraesCS2A02G005900.1 |  | 2758823 | 2760115 | [IPR001128](http://www.ebi.ac.uk/interpro/entry/IPR001128) | Cytochrome P450 |
|  |  |  |  | [IPR002401](http://www.ebi.ac.uk/interpro/entry/IPR002401) | Cytochrome P450, E-class, group I |
|  |  |  |  | [IPR017972](http://www.ebi.ac.uk/interpro/entry/IPR017972) | Cytochrome P450, conserved site |
|  |  |  |  | [IPR036396](http://www.ebi.ac.uk/interpro/entry/IPR036396) | Cytochrome P450 superfamily |
| TraesCS2A02G006000.1 |  | 2791671 | 2794776 | [IPR001906](http://www.ebi.ac.uk/interpro/entry/IPR001906) | Terpene synthase, N-terminal domain |
|  |  |  |  | [IPR005630](http://www.ebi.ac.uk/interpro/entry/IPR005630) | Terpene synthase, metal-binding domain |
|  |  |  |  | [IPR008930](http://www.ebi.ac.uk/interpro/entry/IPR008930) | Terpenoid cyclases/protein prenyltransferase alpha-alpha toroid |
|  |  |  |  | [IPR008949](http://www.ebi.ac.uk/interpro/entry/IPR008949) | Isoprenoid synthase domain superfamily |
|  |  |  |  | [IPR034741](http://www.ebi.ac.uk/interpro/entry/IPR034741) | Terpene cyclase-like 1, C-terminal domain |
|  |  |  |  | [IPR036965](http://www.ebi.ac.uk/interpro/entry/IPR036965) | Terpene synthase, N-terminal domain superfamily |
|  |  |  |  | [IPR044814](http://www.ebi.ac.uk/interpro/entry/IPR044814) | Terpene cyclases, class 1, plant |
|  |  |  |  | [IPR050148](http://www.ebi.ac.uk/interpro/entry/IPR050148) | Terpene synthase-like |
| TraesCS2A02G006100.1 |  | 2817615 | 2820056 | [IPR000719](http://www.ebi.ac.uk/interpro/entry/IPR000719) | Protein kinase domain |
|  |  |  |  | [IPR000858](http://www.ebi.ac.uk/interpro/entry/IPR000858) | S-locus glycoprotein domain |
|  |  |  |  | [IPR001480](http://www.ebi.ac.uk/interpro/entry/IPR001480) | Bulb-type lectin domain |
|  |  |  |  | [IPR003609](http://www.ebi.ac.uk/interpro/entry/IPR003609) | PAN/Apple domain |
|  |  |  |  | [IPR008271](http://www.ebi.ac.uk/interpro/entry/IPR008271) | Serine/threonine-protein kinase, active site |
|  |  |  |  | [IPR011009](http://www.ebi.ac.uk/interpro/entry/IPR011009) | Protein kinase-like domain superfamily |
|  |  |  |  | [IPR017441](http://www.ebi.ac.uk/interpro/entry/IPR017441) | Protein kinase, ATP binding site |
|  |  |  |  | [IPR024171](http://www.ebi.ac.uk/interpro/entry/IPR024171) | S-receptor-like serine/threonine-protein kinase |
|  |  |  |  | [IPR036426](http://www.ebi.ac.uk/interpro/entry/IPR036426) | Bulb-type lectin domain superfamily |
| TraesCS2A02G006200.1 |  | 2839126 | 2854913 | [IPR023213](http://www.ebi.ac.uk/interpro/entry/IPR023213) | Chloramphenicol acetyltransferase-like domain superfamily |
|  |  |  |  | [IPR051283](http://www.ebi.ac.uk/interpro/entry/IPR051283) | Secondary metabolite acyltransferase |
| TraesCS2A02G006300.1 |  | 2860739 | 2862513 | [IPR001128](http://www.ebi.ac.uk/interpro/entry/IPR001128) | Cytochrome P450 |
|  |  |  |  | [IPR002401](http://www.ebi.ac.uk/interpro/entry/IPR002401) | Cytochrome P450, E-class, group I |
|  |  |  |  | [IPR017972](http://www.ebi.ac.uk/interpro/entry/IPR017972) | Cytochrome P450, conserved site |
|  |  |  |  | [IPR036396](http://www.ebi.ac.uk/interpro/entry/IPR036396) | Cytochrome P450 superfamily |
| TraesCS2A02G006400.1 |  | 2876139 | 2877842 | [IPR001128](http://www.ebi.ac.uk/interpro/entry/IPR001128) | Cytochrome P450 |
|  |  |  |  | [IPR002401](http://www.ebi.ac.uk/interpro/entry/IPR002401) | Cytochrome P450, E-class, group I |
|  |  |  |  | [IPR017972](http://www.ebi.ac.uk/interpro/entry/IPR017972) | Cytochrome P450, conserved site |
|  |  |  |  | [IPR036396](http://www.ebi.ac.uk/interpro/entry/IPR036396) | Cytochrome P450 superfamily |
| TraesCS2A02G006500.1 |  | 2880736 | 2885432 | [IPR002182](http://www.ebi.ac.uk/interpro/entry/IPR002182) | NB-ARC |
|  |  |  |  | [IPR027417](http://www.ebi.ac.uk/interpro/entry/IPR027417) | P-loop containing nucleoside triphosphate hydrolase |
|  |  |  |  | [IPR032675](http://www.ebi.ac.uk/interpro/entry/IPR032675) | Leucine-rich repeat domain superfamily |
|  |  |  |  | [IPR036388](http://www.ebi.ac.uk/interpro/entry/IPR036388) | Winged helix-like DNA-binding domain superfamily |
|  |  |  |  | [IPR041118](http://www.ebi.ac.uk/interpro/entry/IPR041118) | Disease resistance, N-terminal |
|  |  |  |  | [IPR042197](http://www.ebi.ac.uk/interpro/entry/IPR042197) | Apoptotic protease-activating factors, helical domain |
|  |  |  |  | [IPR055414](http://www.ebi.ac.uk/interpro/entry/IPR055414) | Disease resistance R13L4/SHOC-2-like, LRR domain |
|  |  | 2892625 | 2893791 | [IPR001810](http://www.ebi.ac.uk/interpro/entry/IPR001810) | F-box domain |
|  |  |  |  | [IPR005174](http://www.ebi.ac.uk/interpro/entry/IPR005174) | KIB1-4, beta-propeller |
|  |  |  |  | [IPR036047](http://www.ebi.ac.uk/interpro/entry/IPR036047) | F-box-like domain superfamily |
| TraesCS2A02G006700.1 |  | 2895357 | 2896658 | [IPR023213](http://www.ebi.ac.uk/interpro/entry/IPR023213) | Chloramphenicol acetyltransferase-like domain superfamily |
|  |  |  |  | [IPR051283](http://www.ebi.ac.uk/interpro/entry/IPR051283) | Secondary metabolite acyltransferase |
| TraesCS2A02G006800.1 |  | 2914803 | 2920979 | [IPR002182](http://www.ebi.ac.uk/interpro/entry/IPR002182) | NB-ARC |
|  |  |  |  | [IPR027417](http://www.ebi.ac.uk/interpro/entry/IPR027417) | P-loop containing nucleoside triphosphate hydrolase |
|  |  |  |  | [IPR032675](http://www.ebi.ac.uk/interpro/entry/IPR032675) | Leucine-rich repeat domain superfamily |
|  |  |  |  | [IPR036388](http://www.ebi.ac.uk/interpro/entry/IPR036388) | Winged helix-like DNA-binding domain superfamily |
|  |  |  |  | [IPR041118](http://www.ebi.ac.uk/interpro/entry/IPR041118) | Disease resistance, N-terminal |
|  |  |  |  | [IPR055414](http://www.ebi.ac.uk/interpro/entry/IPR055414) | Disease resistance R13L4/SHOC-2-like, LRR domain |
| TraesCS2A02G006900.1 |  | 2964420 | 2965424 | [IPR002182](http://www.ebi.ac.uk/interpro/entry/IPR002182) | NB-ARC |
|  |  |  |  | [IPR027417](http://www.ebi.ac.uk/interpro/entry/IPR027417) | P-loop containing nucleoside triphosphate hydrolase |
|  |  |  |  | [IPR041118](http://www.ebi.ac.uk/interpro/entry/IPR041118) | Disease resistance, N-terminal |
| TraesCS2A02G007000.1 |  | 2986007 | 2990362 | [IPR002182](http://www.ebi.ac.uk/interpro/entry/IPR002182) | NB-ARC |
|  |  |  |  | [IPR027417](http://www.ebi.ac.uk/interpro/entry/IPR027417) | P-loop containing nucleoside triphosphate hydrolase |
|  |  |  |  | [IPR032675](http://www.ebi.ac.uk/interpro/entry/IPR032675) | Leucine-rich repeat domain superfamily |
|  |  |  |  | [IPR036388](http://www.ebi.ac.uk/interpro/entry/IPR036388) | Winged helix-like DNA-binding domain superfamily |
|  |  |  |  | [IPR041118](http://www.ebi.ac.uk/interpro/entry/IPR041118) | Disease resistance, N-terminal |
|  |  |  |  | [IPR042197](http://www.ebi.ac.uk/interpro/entry/IPR042197) | Apoptotic protease-activating factors, helical domain |
| TraesCS2A02G007100.1 |  | 3099245 | 3101998 | [IPR000109](http://www.ebi.ac.uk/interpro/entry/IPR000109) | Proton-dependent oligopeptide transporter family |
|  |  |  |  | [IPR036259](http://www.ebi.ac.uk/interpro/entry/IPR036259) | MFS transporter superfamily |
| TraesCS2A02G007200.2 |  | 3106369 | 3111490 |  |  |
| TraesCS2A02G007300.1 |  | 3111972 | 3114111 |  |  |
| TraesCS2A02G007400.1 |  | 3258838 | 3280577 | [IPR002110](http://www.ebi.ac.uk/interpro/entry/IPR002110) | Ankyrin repeat |
|  |  |  |  | [IPR026961](http://www.ebi.ac.uk/interpro/entry/IPR026961) | PGG domain |
|  |  |  |  | [IPR036770](http://www.ebi.ac.uk/interpro/entry/IPR036770) | Ankyrin repeat-containing domain superfamily |
| TraesCS2A02G007500.1 |  | 3333157 | 3336110 | [IPR000109](http://www.ebi.ac.uk/interpro/entry/IPR000109) | Proton-dependent oligopeptide transporter family |
|  |  |  |  | [IPR018456](http://www.ebi.ac.uk/interpro/entry/IPR018456) | PTR2 family proton/oligopeptide symporter, conserved site |
|  |  |  |  | [IPR036259](http://www.ebi.ac.uk/interpro/entry/IPR036259) | MFS transporter superfamily |
|  |  |  |  | [IPR000109](http://www.ebi.ac.uk/interpro/entry/IPR000109) | Proton-dependent oligopeptide transporter family |
|  |  |  |  | [IPR036259](http://www.ebi.ac.uk/interpro/entry/IPR036259) | MFS transporter superfamily |
| TraesCS2A02G007600.1 |  | 3351281 | 3351679 |  |  |
| TraesCS2A02G007700.1 |  | 3400981 | 3406474 | [IPR003854](http://www.ebi.ac.uk/interpro/entry/IPR003854) | Gibberellin regulated protein |
